# NGLY1 knockdown or pharmacological inhibition induces cellular autophagy

**DOI:** 10.1101/2020.12.05.400481

**Authors:** Sarah H Needs, Martin D Bootman, Jeff E Grotzke, Holger B Kramer, Sarah A Allman

**Affiliations:** Reading School of Pharmacy, University of Reading, Whiteknights, Reading, RG6 6AD, UK; School of Life, Health and Chemical Sciences, The Open University, Walton Hall, Milton Keynes, MK7 6AA, UK; Yale University School of Medicine, New Haven, CT 06520, USA; Department of Physiology, Anatomy and Genetics, University of Oxford, Parks Road, Oxford, OX1 3PT, UK; MRC London Institute of Medical Sciences, Hammersmith Hospital Campus, Du Cane Road, London, W12 0NN, UK; Leicester School of Pharmacy, De Montfort University, The Gateway, Leicester, LE1 9BH, UK

**Keywords:** NGLY1, N-Glycanase, Z-VAD-fmk, Q-VD-OPh, caspase inhibition, off-target, autophagy, ERAD, autophagosome proteomics

## Abstract

Pan-caspase inhibitor Z-VAD-fmk acts as an inhibitor of peptide:*N*-glycanase (NGLY1); an endoglycosidase which cleaves *N*-linked glycans from glycoproteins exported from the endoplasmic reticulum during ER-associated degradation (ERAD). Pharmacological *N*-glycanase inhibition by Z-VAD-fmk or siRNA knockdown (KD) induces GFP-LC3 positive puncta in HEK 293 cells. Activation of ER stress markers or reactive oxygen species (ROS) induction are not observed. In NGLY1 inhibition or KD, upregulation of autophagosome formation without impairment of autophagic flux are observed. Enrichment and proteomics analysis of autophagosomes after Z-VAD-fmk treatment or NGLY1 KD reveals comparable autophagosomal protein content. Upregulation of autophagy represents a cellular adaptation to NGLY1 inhibition or KD, and ATG13-deficient mouse embryonic fibroblasts (MEFs) show reduced viability under these conditions. In contrast, treatment with pan-caspase inhibitor, Q-VD-OPh does not induce cellular autophagy. Therefore, experiments with Z-VAD-fmk are complicated by the effects of NGLY1 inhibition and Q-VD-OPh represents an alternative caspase inhibitor free from this limitation.

## Introduction

The peptide fluoromethyl ketone inhibitor Z-VAD-fmk has been applied extensively to the study of caspases and the central role of this important group of cysteine proteases in apoptosis. The widespread use of this inhibitor is likely due to its effective inhibition of a range of caspases and its extensive application for over more than 20 years in biological research. Utilization as a tool in cellular research continues to date, despite the recognition that Z-VAD-fmk also interacts with several off-targets. These off-targets include other cysteine proteases, such as the cathepsins (Schotte et al., 1999) and calpains (Waterhouse et al., 1998), but also the amidase peptide:*N*-Glycanase 1 (NGLY1) (Misaghi et al., 2004). Additionally, it has been shown that both Z-VAD-fmk and Z-IETD-fmk, a related peptide fluoromethyl ketone (fmk) inhibitor, are capable of inhibiting rhinoviral proteinases and thereby reduce viral replication in cultured cells (Deszcz et al., 2004). The selectivity profile of Z-VAD-fmk across human caspases has been characterized in detail (Garcia-Calvo et al., 1998). Furthermore, quantitative comparisons of inhibition profiles between Z-VAD-fmk and Q-VD-OPh across caspases and cathepsins has shown that the latter represents a more potent and more selective second-generation inhibitor (Caserta et al., 2003; Chauvier et al., 2007). Despite this, Z-VAD-fmk is still in recent and current use in a large number of studies including for the induction of necroptosis or continued use for the inhibition of caspase-mediated apoptosis. While inhibition of cellular caspases leads to suppression of apoptosis, it has been shown that Z-VAD-fmk treatment can induce other forms of programmed cell death, such as autophagic cell death in L929 fibrosarcoma cells (Chen et al., 2011). The authors showed that cellular autophagy, including autophagosome and autolysosome formation, was central in this pathway to cell death and implicated inactivated caspase 8 and increased ROS production in the process. Disparate cellular effects of different caspase inhibitors have also been documented in the literature. Investigations with the inhibitors Z-VAD-fmk, Boc-D-fmk and Q-VD-OPh in L929 cells revealed that the two fmk-based inhibitors induced necroptosis, while Q-VD-OPh did not (Wu et al., 2011). The investigators went on to show that autocrine production of TNFα was involved in this process and was mediated by the PKC-MAPKs-AP1 pathway. Furthermore, they demonstrated that the NF-kB pathway exerts a protective function against necroptosis. Despite such disparate cellular outcomes for different caspase inhibitors, the precise mechanistic reasons for varying outcomes with different inhibitors are often unclear. The fmk pharmacophore has been implicated in potential undesired effects due to metabolic turnover into toxic fluoroacetate (Eichhold et al., 1997) which, following conversion to fluorocitrate, is capable of inactivating aconitase within the citric acid cycle (Morrison and Peters, 1954). It should be noted however that significant differences in selectivity between fmk-based and alternative inhibitors itself may explain differences in engagement of cellular off-targets.

The primary biochemical function of the known Z-VAD-fmk off-target NGLY1 is the enzymatic removal of *N*-linked glycans from glycoprotein substrates (Suzuki et al., 2016; Suzuki et al., 1993) and its involvement in the Endoplasmic Reticulum-Associated Degradation (ERAD) pathway has been clearly demonstrated (Grotzke et al., 2013; Suzuki et al., 2015). Furthermore, NGLY1 deficiency is a rare genetic disorder with a complex clinical presentation which often includes neurological symptoms and developmental delay (Enns et al., 2014; He et al., 2015). Recent findings have demonstrated novel cellular functions of NGLY1 in the regulation of aquaporins independent of its enzymatic activity (Tambe et al., 2019) and *N*-glycanase dependent sequence editing of Nrf-1, leading to the induction of gene expression of proteasomal subunits (Lehrbach et al., 2019; Tomlin et al., 2017; Yang et al., 2018). Recently it has also been shown that knockdown (KD) of NGLY1 in K562 human myelogenous leukemia cells increases sensitivity towards proteasome inhibition and induces complex changes in transcript and protein regulation (Mueller et al., 2020). Despite these important findings there is still limited mechanistic information on the effects of NGLY1 inhibition or disruption in mammalian cells.

The aim of this study is to delineate to which extent the cellular effects observed upon treatment with Z-VAD-fmk are mediated by the inhibition of its off-target NGLY1.

## Results

### Pan-caspase inhibitor Z-VAD-fmk, but not Q-VD-OPh inhibits *N*-glycanase activity of NGLY1

We were interested in investigating the cellular effects of inhibition of NGLY1 by Z-VAD-fmk, a known off-target for this pan-caspase inhibitor. siRNA-mediated gene knock-down (KD) in HEK 293 cells leads to a significant reduction in NGLY1 transcript levels (Fig. 1a) while cell viability remains unaffected (Fig. 1b). In order to assess remaining enzymatic *N*-glycanase activity in NGLY1 KD cells, a deglycosylation-dependent Venus (ddVenus) assay (Grotzke et al., 2013) was employed. This demonstrated a significant reduction of *N*-glycanase activity in NGLY1 KD cells by 68% at 72 h post-transfection (Fig.1c). A similar significant reduction of 72% of *N*-glycanase activity was observed with the same ddVenus assay following treatment of HEK 293 cells with 50 μM Z-VAD-fmk for 24 h (Fig 1d). Conversely, HEK 293 cells treated with 50 μM Q-VD-OPh for the same time period showed no significant alteration of fluorescence intensity in the ddVenus assay, indicating that cellular *N*-glycanase activity remained unaffected (Fig 1d).

**Figure 1.**
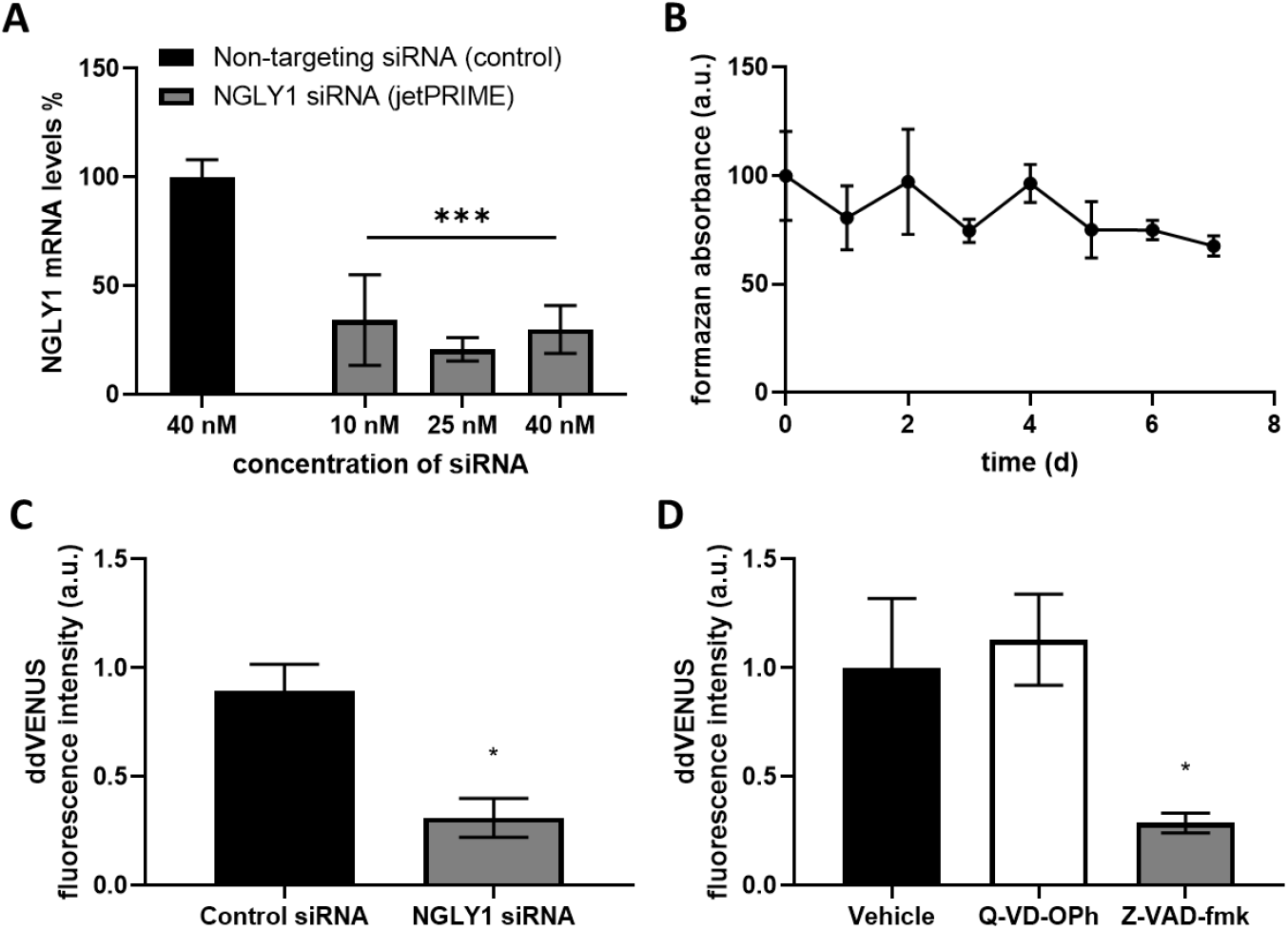
*N*-glycanase siRNA knockdown and Z-VAD-fmk treatment reduce the amount of cellular deglycosylation. (A) Transfection of NGLY1 siRNA in HEK 293 cells results in significant reduction of NGLY1 mRNA levels (compared with non-targeting control). Optimisation of transfection of NGLY1 siRNA in HEK 293 cells. Data represents the percentage mRNA knockdown at different concentrations of targeting siRNA (10, 25 and 40 nM using jetPRIME™ as a transfection reagent) compared to a non-targeting control siRNA. Two-way ANOVA, Dunnett’s post hoc compared to nontargeting siRNA. ** p < 0.01, *** p < 0.001, Error bars ± SEM, n=3. (B) Transfection of NGLY1 siRNA in HEK 293 cells does not cause significant decrease in cellular viability. MTT cell viability assay. HEK 293 cells transfected with 25 nM NGLY1 siRNA over 7 days. Data normalised to non-targeting control of the same time and Day 0 represented as 100 %. One-way ANOVA, followed by a Dunnett’s post hoc test against the vehicle. HEK 293 cells were treated with either Q-VD-OPh or Z-VAD-fmk (50 μM) for 24 h (C) or transfected with NGLY1 siRNA or non-targeting siRNA (25 nM) with jetPRIME^®^ for 3 d (D) followed by transfection with ddVENUS construct with jetPEI^®^. After 72 h cells were treated with MG132 (8 μM) for 6 h and fluorescence intensity was analysed by flow cytometry. The median ddVENUS fluorescence was calculated and normalised to the vehicle. n=3, error bars indicate ± SEM. * indicates p < 0.05.

### Z-VAD-fmk treatment induces cellular autophagy via NGLY1 inhibition in HEK 293 cells

Given the different effects of Z-VAD-fmk and Q-VD-OPh with respect to inhibition of cellular *N*-glycanase activity, we were interested in comparing the cellular effects of the two inhibitors more broadly. Previous reports have implicated an involvement of autophagy in necrotic cell death which can be observed in some cell lines treated with Z-VAD-fmk (Chen et al., 2011; Wu et al., 2014; Wu et al., 2011). Treatment of stably transfected GFP-LC3 HEK 293 cells with Z-VAD-fmk (50 μM), Q-VD-OPh (50 μM) or vehicle control for 24, 48 or 72 h showed a significant increase in the number of GFP-LC3 puncta per cell at the 72 h time point in the Z-VAD-fmk treated cells (Fig 2a, 2b).

**Figure 2.**
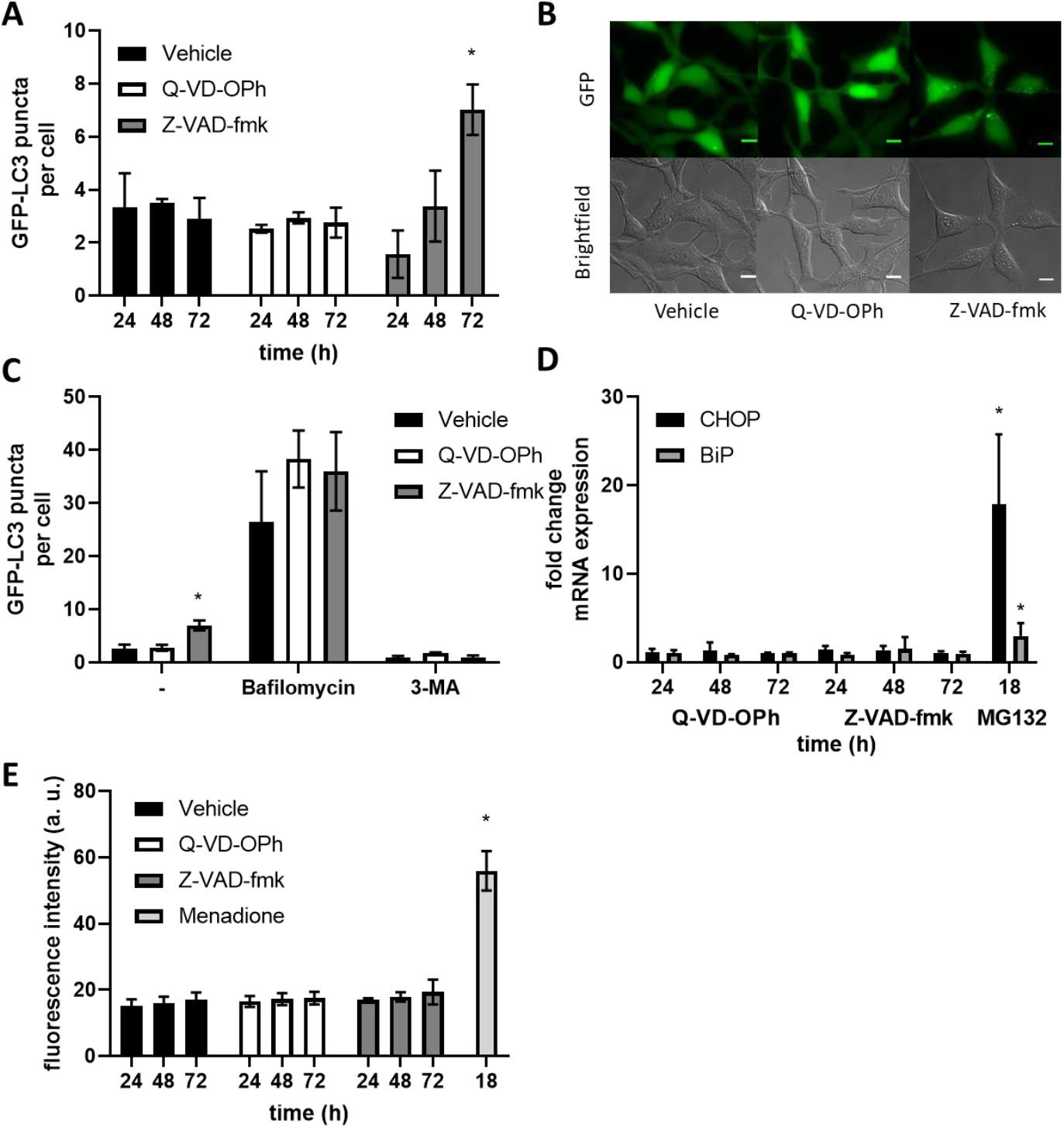
Z-VAD-fmk, but not Q-VD-OPh induces autophagy. (A) GFP-LC3 puncta per cell after treatment with Z-VAD-fmk or Q-VD-OPh (50 μM, 24-72 h). (B) Representative images of treatment of Z-VAD-fmk or Q-VD-OPh after 72 h. Scale bar indicates 10 μm. (C) GFP LC3 puncta per cell after 72 h treatment with Z-VAD-fmk or Q-VD-OPh followed by Bafilomycin A1 treatment (1 h, 100 nM) or 3-MA (1 h, 5 mM). Two-way ANOVA, Tukey’s post hoc. (D) rt-qPCR analysis of CHOP and BiP in HEK 293 cells treated with Z-VAD-fmk or Q-VD-OPh (50 μM, 24-72 h). mRNA levels normalised to GAPDH and a vehicle control using ΔΔCt method. Two-way ANOVA, Tukey’s post hoc within each mRNA target. (E) HEK 293 cells treated with Z-VAD-fmk or Q-VD-OPh (50 μM, 24-72 h). Cells were stained with ROS Brite 570 (5 μM, 0.5 h). Two-way ANOVA, Tukey’s post hoc, error bars ± SEM, n=3, * p < 0.05.

### Lack of ER stress or Redox imbalance in cells subjected to Z-VAD-fmk, Q-VD-OPh or NGLY1 KD

As autophagy activation has been reported following ER stress, potential induction of the endoplasmic reticulum (ER) stress markers C/EBP-homologous protein (CHOP) and Binding-immunoglobulin protein (BiP) was monitored using quantitative real-time polymerase chain reaction (qPCR) (Fig 2d). No induction of ER stress markers was observed for either Z-VAD-fmk or Q-VD-OPh treatment (each at 50 μM) at the time points tested (24, 48 and 72 h), while the positive control treatment with proteasome inhibitor MG132 (5 μM, 18 h) showed robust induction of both ER stress markers. We also explored the possible effects on cellular redox homeostasis and production of reactive oxygen species (ROS). To this end, we stained HEK 293 cells with ROS Brite 570 following treatment with either vehicle control, Z-VAD-fmk or Q-VD-OPh (50 μM each). This showed that across treatment periods of 24-72 h, no significant induction of cellular ROS could be observed. As a positive control, menadione treatment (5 μM, 18 h) showed a significant increase of the measured ROS Brite fluorescence (Fig 2e). In order to confirm whether the induction of autophagosome formation, as shown by induction of LC3 puncta, can be explained by the inhibition of NGLY1, we carried out a comparable experiment with siRNA mediated NGLY1 KD (Fig. 3a, 3b). A significant increase in GFP-LC3 puncta was observed at 5 days post-transfection when comparing NGLY1 KD to the non-targeting siRNA control. As with the inhibitor treatment experiments (Fig 2d, 2e) we investigated induction of ER stress markers CHOP and BiP, and cellular ROS in this system. We found no significant differences between non-targeting siRNA control and NGLY1 KD cells (Fig 3d, 3e) and thereby no evidence for induction of ER stress or disruption of cellular redox balance.

**Figure 3.**
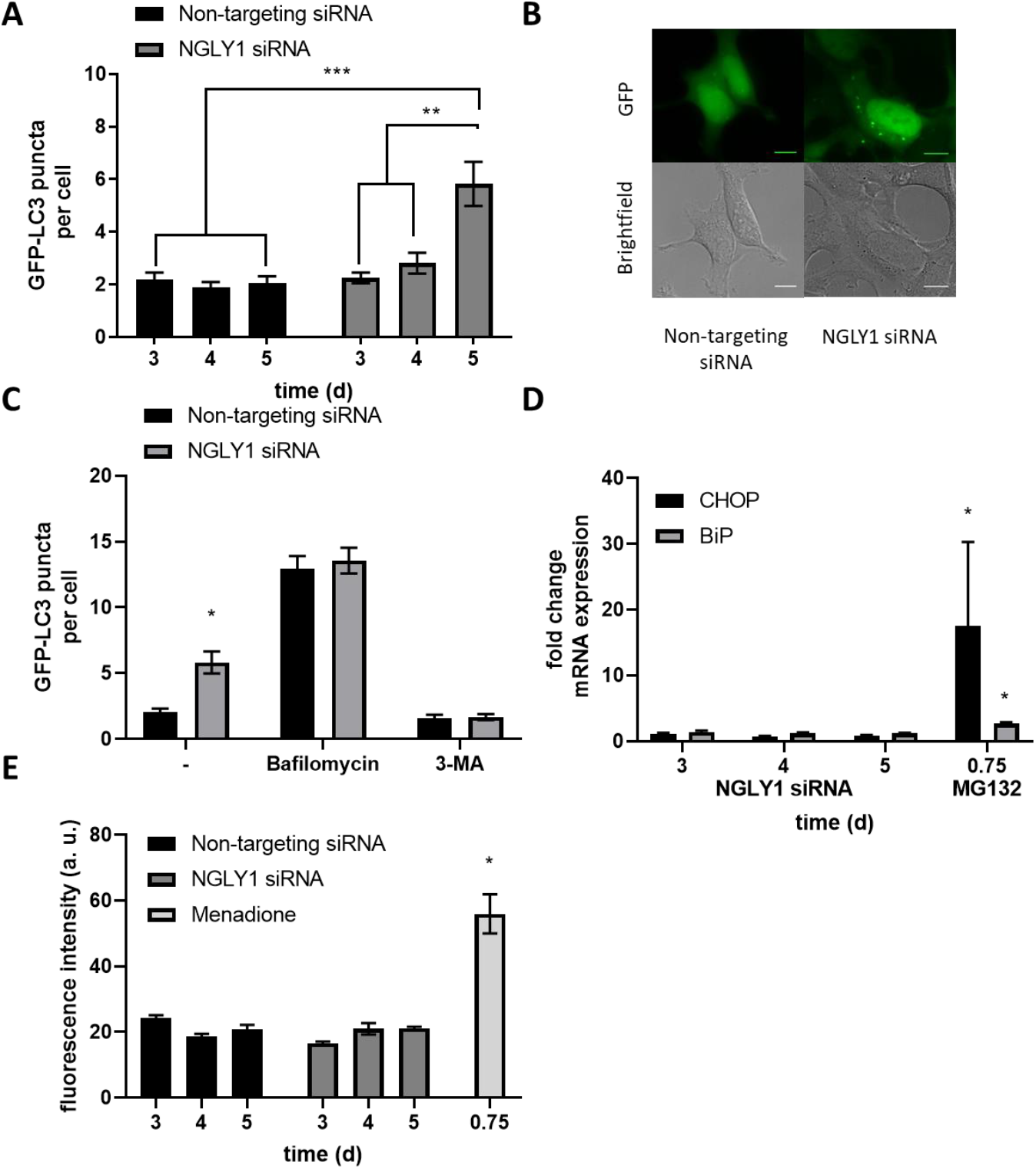
siRNA knockdown of *N*-glycanase induces autophagy. (A) GFP-LC3 puncta are increased after 5 d post-transfection with NGLY1 siRNA (25 nM). Quantitation of the average GFP-LC3 puncta per cell, minimum of three images per coverslip with three biological replicates. Error bars indicate ± SEM, ** p < 0.01, *** p < 0.001, two-way ANOVA, Tukey’s post hoc. (B) Representative images of HEK 293 cells transfected with NGLY1 siRNA or nontargeting siRNA after 5 days. Scale bar indicates 10 μm. (C) GFP-LC3 puncta per cell after transfection with NGLY1 siRNA or non-targeting siRNA (25 nM, 5 d) with Bafilomycin A1 (100 nM, 1 h) or 3-MA (5 mM, 1 h). (D) rt-qPCR anlaysis of CHOP and BiP in HEK 293 cells transfected with NGLY1 siRNA (25 nM, 3-5 days), mRNA levels normalised to GAPDH and a non-targeting siRNA control using ΔΔCt method. Two-way ANOVA, Tukey’s post hoc within each mRNA target. (E) HEK 293 cells transfected with NGLY1 siRNA or non-targeting siRNA (25 nM, 3-5 d) and incubated with ROS Brite™ 570 (5 μM, 0.5 h). Two-way ANOVA, Tukey’s post hoc. * p < 0.05, Error bars ± SEM, n=3.

### Cellular viability upon Z-VAD-fmk treatment is reduced in ATG13 KO compared to WT MEF cells

We also performed MTT assays of wild type mouse embryonic fibroblasts (WT MEFs) and ATG13 knock-out (KO) MEF cells treated with either Z-VAD-fmk or Q-VD-OPh. ATG13 is essential for cellular autophagy and KO cells have strongly diminished ability to induce autophagosome formation (Kaizuka and Mizushima, 2016). Having confirmed ATG13 deficiency in the KO cells by determining transcript levels by qPCR (Fig S2a), we compared the effects of inhibitor treatment of WT and ATG13 KO MEF cells (Fig S2b,c). Upon Z-VAD-fmk treatment a significant difference in formazan absorbance was observed at inhibitor concentrations of 100 and 200 μM (Fig S2b). In contrast, no significant difference between WT and ATG13 KO MEFs was observed for Q-VD-OPh at all concentrations tested (Fig S2c).

### Induction of autophagosome formation rather than disruption of autophagic flux is responsible for NGLY1-mediated increase in cellular autophagy

A further experiment was carried out to assess whether the observed induction of autophagosome formation in Z-VAD-fmk treatment was caused by a disruption of autophagic flux or due to genuine upregulation of cellular autophagosome formation. Treatment of stably transfected GFP-LC3 HEK 293 cells with Z-VAD-fmk or Q-VD-OPh for 72 h was followed by inhibition with Bafilomycin or 3-Methyladenine (3-MA) for 1 h (Fig 2c). This showed no evidence for disruption of cellular autophagic flux. Similarly, treatment of stably transfected GFP-LC3 HEK 293 cells with Bafilomycin or 3-Methyladenine (3-MA) following transfection with either non-targeting control siRNA or NGLY1 KD for 5 days was investigated (Fig 3c). While the previously observed induction of autophagosome formation in NGLY1 KD was recapitulated, there was no significant difference between control siRNA or NGLY1 KD for both Bafilomycin and 3-MA treatment (Fig 3c). This indicates that cellular autophagic flux is not affected in NGLY1 KD and a genuine increase in autophagosome formation is responsible for the observed elevation of GFP-LC3 puncta per cell.

### Intracellular Ca2+ handling is unaffected by treatment with Z-VAD-fmk, Q-VD-OPh or NGLY1 KD

In order to assess whether intracellular Ca^2+^ signaling is affected by Z-VAD-fmk or Q-VD-OPh inhibitor treatment, we studied whether release of Ca^2+^ from intracellular stores is affected under these conditions. HEK 293 cells were loaded with Fura-2 AM dye (1 μM) following inhibitor treatment (vehicle control, Z-VAD-fmk or Q-VD-OPh) and then Ca^2+^ release stimulated with Thapsigargin (1 μM). Fluorescence imaging was carried out with excitation at 340nm and 380nm wavelengths allowing ratiometric determination of free intracellular Ca^2+^ concentrations. These measurements indicated no significant difference in Ca^2+^ release from intracellular stores following inhibitor treatments with either Z-VAD-fmk or Q-VD-OPh (50 μM each, 24-72 h) relative to vehicle control (Fig S3a-f). Comparable experiments were also conducted following NGLY1 KD or treatment with non-targeting control siRNA. Again, no significant difference in Ca^2+^ release from intracellular stores was observed at any of the time points (3-5 d) tested (Fig S4a-e).

### Autophagosome immunoprecipitation and mass spectrometry-based proteomics identifies autophagosomal protein content following Z-VAD-fmk treatment or NGLY1 KD

In order to investigate autophagosomal proteins at the molecular level, we performed autophagosome proteomics analysis of stably transfected GFP-LC3 HEK 293 cells which had undergone either Z-VAD-fmk treatment or NGLY1 KD. Autophagosomes were enriched by centrifugation followed by anti-GFP immunoprecipitation (IP) and corresponding negative control IPs were carried out in parallel. All experiments were conducted as biological triplicates. Sample processing and analysis was carried out by trypsin digestion and liquid chromatography-tandem mass spectrometry (LC-MS/MS). Raw data were then processed and analyzed in MaxQuant (Cox and Mann, 2008) using the Label-free quantification (LFQ) algorithm (Cox et al., 2014) and statistical analysis and data visualization in Perseus (Tyanova et al., 2016). After removal of known contaminant proteins this led to the identification 1,011 protein hits at a False Discovery Rate (FDR) of 1%. Filtering to obtain only proteins which were quantified in at least two out of three replicates per experimental group retained 915 protein hits.

The label-free quantitative proteomics data shows enrichment of a subset of autophagy-related proteins (Table 1) in autophagosomes isolated from Z-VAD-fmk treatment or NGLY1 KD, relative to the corresponding control IPs. The observation of multiple autophagy related proteins validates the experimental workflow and successful enrichment of autophagosomes. Strong enrichment is also shown in volcano plots comparing the GFP IPs to their respective negative control IPs for Z-VAD-fmk treatment or NGLY1 KD (Fig. 4). Among the notable proteins which were found to be significantly enriched in the autophagosome IPs in both inhibitor treatment and NGLY1 KD were the bait protein MAP1LC3B, several known interactors thereof (MAP1A, MAP1B, FYCO1, SQSTM1), several autophagy related proteins (ATG3, ATG4B, ATG7), other known autophagosomal proteins (RAB1B, RAB7A, SH3GLB1) as well as known regulators of autophagy (HDAC6, HDAC10, MTDH). Direct comparison of the two autophagosome IPs from inhibitor treatment and NGLY1 siRNA KD was also performed by volcano plot (Fig S5b) indicating no significant changes between the two IPs. A large degree of similarity between the two IPs is also evident in the heatmap with hierarchical clustering analysis (HCA) (Fig. 5). Filtering for ANOVA significant (FDR 0.05) proteins prior to HCA yields 369 significantly altered protein hits. These are subdivided into two clusters of proteins enriched in the autophagosome IPs (cluster I, 228 protein hits) and proteins enriched in the negative control IPs (cluster II, 141 protein hits). Investigation of the protein hits represented in clusters I and II by gene ontology (GO) analysis highlights biological processes (GOBP), cellular compartments (GOCC) and molecular functions (GOMF) which are overrepresented in the respective cluster (Supplementary Tables ST1 and ST2). For proteins enriched in the autophagosome IPs the GOBP terms which are overrepresented include protein translation (translation, translational elongation, translational termination, translational initiation), protein localization and targeting (SRP-dependent cotranslational protein targeting to membrane, protein targeting to ER, establishment of protein localization to organelle, establishment of protein localization in endoplasmic reticulum, protein targeting to membrane, cotranslational protein targeting to membrane, establishment of localization), RNA degradation (nuclear-transcribed mRNA catabolic process - nonsense-mediated decay, mRNA catabolic process, RNA catabolic process nuclear-transcribed mRNA catabolic process) and protein complex disassembly (protein complex disassembly, cellular component disassembly, macromolecular complex disassembly, cellular component disassembly at cellular level, cellular macromolecular complex disassembly). GOCC and GOMF terms overrepresented in the autophagosome IPs are largely focused on ribosomes (GOCC: cytosolic large ribosomal subunit, large ribosomal subunit; GOMF: structural constituent of ribosome). These results confirm successful enrichment of autophagosomes following Z-VAD-fmk treatment or NGLY1 KD, identification of similar autophagosomal protein composition for both conditions and the identification of autophagosomal cargo proteins and the corresponding biological processes represented.

**Table 1.**
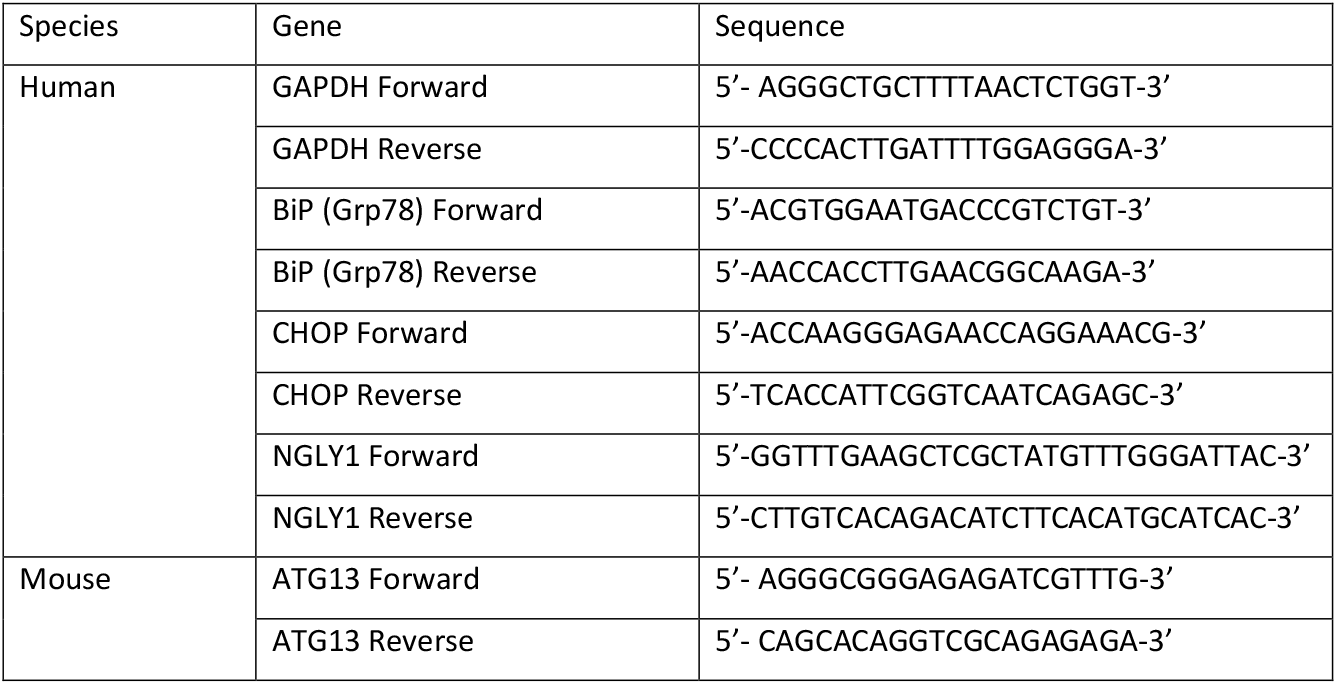
rt-qPCR Primer Sequences

**Figure 4.**
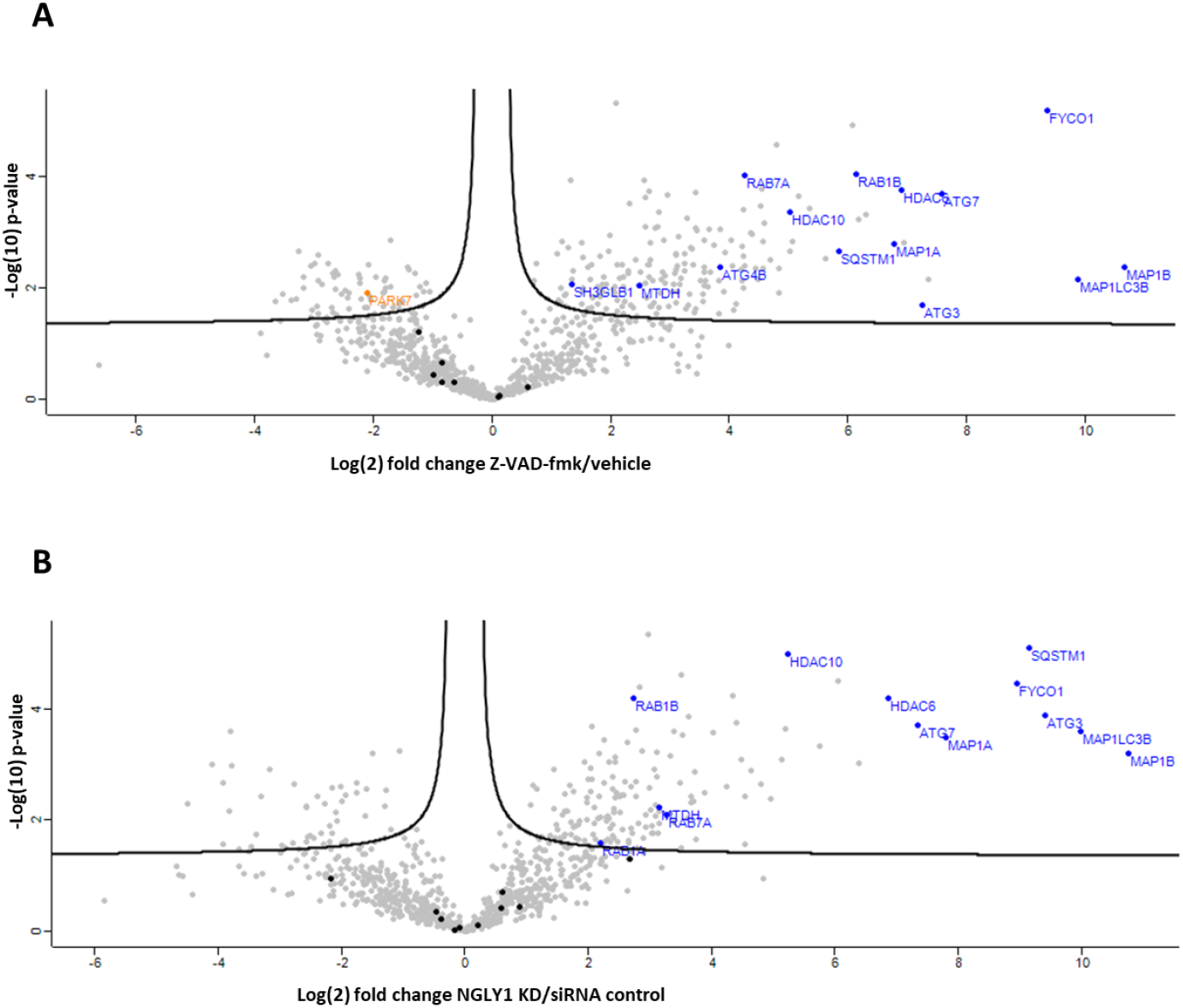
Label-free proteomics analysis of autophagosomes isolated from HEK 293 cells treated with Z-VAD-fmk or NGLY1 KD. Volcano plots identifying significantly enriched and depleted proteins (FDR: 0.05, s0: 0.1) in autophagosomes enriched following (A) Z-VAD-fmk treatment or (B) NGLY1 KD relative to negative control IPs. Autophagy-related proteins are highlighted in blue (significantly enriched), black (not significantly altered) or orange (significantly depleted).

**Figure 5.**
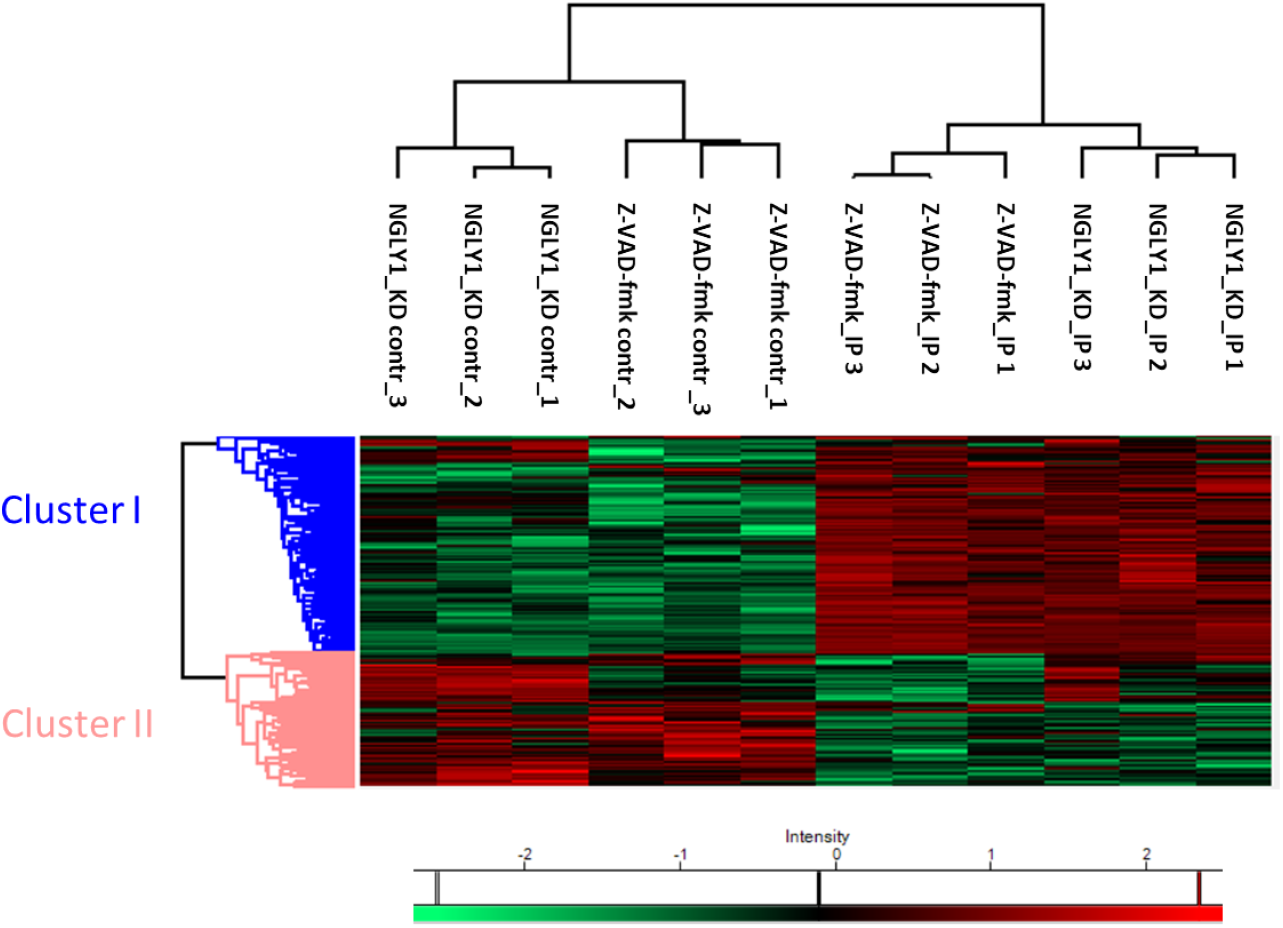
Label-free quantitative proteomics identifies significantly altered protein hits in autophagosome IPs following Z-VAD-fmk treatment or NGLY1 KD. Heatmap and hierarchical clustering analysis (HCA) of significantly changing (One-way ANOVA, FDR: 0.05) protein hits quantified following autophagosome IP or negative control IP and LC-MS/MS analysis. Log(2)-transformed Label-free Quantification (LFQ) intensities after Z-scoring and filtering to retain only ANOVA significant hits. Two clusters represent protein hits enriched in in autophagosome IPs (cluster I) and protein hits enriched negative control IPs (cluster II).

## Discussion

Interest in NGLY1 has been focused on its central role in rare genetic disorder NGLY1 deficiency and in its importance towards cellular glycoprotein turnover. NGLY1 is also a known off-target of the pan-caspase inhibitor Z-VAD-fmk and our results show that siRNA-mediated knockdown of NGLY1 achieves a comparable reduction in cellular peptide:*N*-glycanase activity as that observed with Z-VAD-fmk treatment (24h, 50 μM) (Fig 1c,d). In contrast, treatment with the alternative caspase inhibitor Q-VD-OPh does not affect cellular peptide:N-glycanase activity in HEK 293 cells (Fig 1d), a result which is in agreement with previous studies on *S.cerevisiae* PNG1 (Misaghi et al., 2004). A broader investigation of cellular responses to inhibitor treatment showed no evidence of induction of ER stress markers or ROS production upon Z-VAD-fmk or Q-VD-OPh treatment in HEK 293 cells (Fig 2d,e). However, induction of cellular autophagosome formation (determined by quantification of GFP-LC3 puncta) was observed following treatment with Z-VAD-fmk at 72 h (50 μM), but not with Q-VD-OPh treatment in stably transfected GFP-LC3 HEK 293 cells (Fig 2a,b). Induction of autophagosome formation was also observed with siRNA-mediated KD of NGLY1 at 5 days post-transfection, but not with the scrambled control siRNA (Fig 3a,b). These results strongly suggest that induction of autophagosomes in Z-VAD-fmk treatment is related to NGLY1 inhibition and not to caspase inhibition or other potential off-targets. As for the inhibitor treatment experiments, ER stress markers or ROS production were determined and found not to be altered in NGLY1 KD relative to control (Fig 3d,e). This result agrees with previous studies carried out on NGLY1 KD in *Drosophila melanogaster* (Owings et al., 2018) and in NGLY1 knockout (KO) rats (Asahina et al., 2020) where no evidence of induction of ER stress was found. In addition, the rat model showed evidence for the accumulation of polyubiquitinated proteins in neurons from five-week-old NGLY1 KO rats (Asahina et al., 2020). The study in the *Drosophila* model also showed evidence for cnc (fly ortholog of NRF1) dysfunction by transcriptome analysis among downregulated genes (such as those encoding for proteasomal subunits) and an upregulation of the heat shock response (Owings et al., 2018). Similar downregulation at the transcriptional level of proteasomal subunits was also observed following NGLY1 KD in human K562 cells and was interpreted as related to reduced post-translational processing of NFE2L1 to the mature transcription factor NRF1 by NGLY1 (Mueller et al., 2020). Consistent with these observations it was found that NGLY1 disruption sensitizes towards proteasome inhibition by bortezomib in human cells (Mueller et al., 2020) and *Drosophila* larvae (Rodriguez et al., 2018). The NGLY1 KO mouse model showed the importance of the genetic background towards embryonic lethality and demonstrated that concomitant KO of the Engase gene partially rescued embryonic lethality in C57BL/6 mice (Fujihira et al., 2017). It has been suggested that this observation may be explained by a detrimental effect of *N*-GlcNAc modified proteins formed by ENGase-mediated processing of *N*-linked glycoproteins in the absence of functional NGLY1 (Maynard et al., 2020).

It is noteworthy that an induction of autophagy upon Z-VAD-fmk treatment has been observed previously in experiments with L929 cells where autophagic cell death is induced (Chen et al., 2011; Wu et al., 2014; Wu et al., 2011) and patient-derived tenofibroblasts where autophagy induction is observed in the presence or absence of hydrogen peroxide co-treatment (Kim et al., 2014). This suggests that Z-VAD-fmk mediated autophagy induction is not restricted to a narrow range of cells or cell lines. However, to the best of our knowledge it has not been demonstrated before that this cellular effect is mediated by inhibition of NGLY1. In order to pinpoint the origin of the increase in levels of cellular autophagosomes, we compared blockage of lysosomal degradation with bafilomycin or inhibition of autophagosome biogenesis using 3-Methyladenine (3-MA) for each of vehicle, Z-VAD-fmk or Q-VD-OPh treatments at the 72 h timepoint (Fig 2c). The observed increases (bafilomycin) and decreases (3-MA) in autophagosome content for all three conditions indicate that autophagic flux is not impaired in this cell model. This strongly suggests that the observed increased number of GFP-LC3 puncta is the result of genuine induction of autophagosome formation and not a defect of autophagic flux. Importantly, corresponding experiments were also carried out with control siRNA and NGLY1 KD cells at 5 days post-transfection, similarly showing no evidence of impairment in autophagic flux in cells deficient in NGLY1 (Fig 3c). While an increase in autophagosome formation is a consequence of both Z-VAD-fmk treatment and NGLY1 KD, autophagy also represents an adaptation to reduced cellular *N*-glycanase activity. Investigations using ATG13 KO MEF cells show that these ATG13 deficient cells (Fig S2a) are less tolerant towards treatment with Z-VAD-fmk. Cellular viability upon treatment with Z-VAD-fmk, as assessed by MTT reduction, was significantly reduced in ATG13 KO MEF cells when compared to control WT MEF cells (Fig S2b). In contrast to this, no such difference between ATG13 KO and WT MEF cells could be observed for treatment with Q-VD-OPh across the concentration range of 0-200 μM (Fig S2c). It has been shown previously that Resveratrol-induced autophagy, which is triggered via a non-canonical pathway, involves inositol triphosphate receptors and cytosolic Ca^2+^ in HEK293 cells (Luyten et al., 2017). Similar observations have also been made previously in the context of Resveratrol-induced autophagic cell death in A549 cells (Zhang et al., 2013). Therefore, we decided to investigate whether alterations in intracellular Ca^2+^ signaling are observed in Z-VAD-fmk or Q-VD-OPh inhibitor treatment or following NGLY1 KD. A further motivation for investigating cellular Ca^2+^ signaling was the observation that NGLY1 deficiency leads to impairment in mitochondrial function (Kong et al., 2018; Yang et al., 2018) which could conceivably contribute to alterations in intracellular Ca^2+^ handling. However our results (Fig S3a-f, S4a-e) indicate that neither treatment with Z-VAD-fmk or Q-VD-OPh nor NGLY1 KD significantly affect release of Ca^2+^ from intracellular stores by Thapsigargin stimulation.

We carried out autophagosome enrichment by a two-step process of centrifugation followed by IP of GFP-LC3, in order to investigate autophagosomal protein content. LC3 specifically associates with autophagosomes following post-translational processing to LC3-II and represents the primary autophagosomal marker (Kabeya et al., 2000). The enriched autophagosomes were then processed by lysis, protein denaturation and trypsin digestion for LC-MS/MS and quantitative label-free proteomics analysis (Fig S5a). By conducting and processing negative control precipitations in parallel utilising agarose beads, we assessed enrichment relative to control in order to identify autophagosome-associated proteins and autophagosomal cargo. These experiments were carried out for both Z-VAD-fmk treatment and NGLY1 KD and were conducted in biological triplicates. Following data analysis and processing using previously established methods, enrichment of proteins was visualised using volcano plots (Fig 4a,b). On the plots 22 autophagy-related proteins (Table 1) which were quantified in the analysis were highlighted and 14 of these were found to be significantly enriched (FDR: 0.05, s0: 0.1) following Z-VAD-fmk treatment and 13 significantly enriched in autophagosome IPs after NGLY1 KD (Table 1, Fig 4a,b). Importantly, this included the bait protein LC3 and the known LC3-interacting proteins MAP1A (Mann and Hammarback, 1994), MAP1B (Harrison et al., 2008; Mann and Hammarback, 1994), FYCO1 (Pankiv et al., 2010) and SQSTM1 (Tang et al., 2013) all of which were significant and highly enriched. Significant enrichment in both IP experiments was also observed for the autophagy proteins ATG3 and ATG7. Similarly, the regulator of autophagosome maturation HDAC6 (Lee et al., 2010), the positive autophagy regulator HDAC10 (Oehme et al., 2013) and autophagy inducer MTDH (Bhutia et al., 2010) were found to be enriched and significant in both experiments. Direct comparison of the two autophagosome IPs following Z-VAD-fmk treatment or NGLY1 KD by volcano plot showed that there were no significantly altered protein hits between these samples (FDR: 0.05, s0: 0.1), suggesting that both triggers of autophagy lead to comparable cellular autophagic responses and autophagosomal protein cargo (Fig S5b). The entire proteomics dataset was also displayed as a heatmap of protein abundance, filtered for ANOVA significant protein hits (FDR 0.05) and analysed by hierarchical clustering analysis (HCA) (Fig 5). This approach further confirmed the similarity of proteins significantly enriched in the autophagosome IP experiments (cluster I, representing 228 protein hits). Gene Ontology (GO) analysis for biological process terms showed enrichment of protein translation, protein localization and targeting, mRNA degradation and protein complex disassembly. GO terms for cellular compartment and molecular function showed that ribosomal proteins were overrepresented. Given the generation of autophagosomes from phagophores formed from the ER, it is plausible that a range of ER-resident, ribosomal and membrane proteins are overrepresented in autophagosomes.

Previous work which investigated autophagosome isolation from HEK 293 cells, followed by 2D gel electrophoresis, in-gel digestion and analysis by MALDI-TOF mass spectrometry identified 101 proteins (Gao et al., 2010). Of this list of putative autophagosomal proteins, we also identified 23 proteins in our data of which 9 (Z-VAD-fmk) and 7 (NGLY1 KD) were found to be significantly enriched in our analyses (Fig S6a,b). It should be noted that in addition to the different processing and analysis approaches, autophagy was induced by calcium phosphate precipitate in this previous study (Gao et al., 2010). One further comparison was performed to a list of 94 autophagosomal proteins enriched from MCF-7 cells which were determined to localise to autophagosomes independent of the trigger of cellular autophagy (Dengjel et al., 2012). Here a greater overlap of 37 out of the 94 proteins which were also identified in our study was observed. However significant enrichment was seen for only 8 (Z-VAD-fmk) and 6 (NGLY1 KD) of these 37 proteins with most of the remainder of proteins identified, but found not to be significantly altered in our experiments (Fig S7a,b). This suggests that the more comparable experimental approaches (analysis by LC-MS/MS and control for FDR using target-decoy database search approach) enabled a greater overlap of the datasets. However other confounding factors, such as the expected biological differences between MCF-7, as a cancer cell line, as opposed to the HEK 293 non-malignant cells utilised in our studies, result in a number of quantitative differences in autophagosome formation and maturation. These proteomic investigations reveal similar autophagosomal protein composition following either Z-VAD-fmk treatment or NGLY1 KD in HEK 293 cells and extend our understanding of factors which are enriched in cellular autophagosomes following the triggering of cellular autophagy by impairment or inhibition of NGLY1.

In conclusion we have shown that cellular treatment with the pan-caspase inhibitor Z-VAD-fmk leads to an induction of autophagy which is mediated by the inhibition of NGLY1. No such increase in cellular autophagy is observed with the alternative caspase inhibitor Q-VD-OPh. The increase in GFP-LC3 puncta per cell upon Z-VAD-fmk treatment or NGLY1 KD occurs due to an upregulation in autophagosome formation and no disruption of autophagic flux can be observed. Isolation of autophagosomes by IP, following Z-VAD-fmk treatment or NGLY1 KD, and mass spectrometry-based proteomics analysis identifies a similar range of autophagosomal proteins and protein cargo. Analysis of GO terms for the autophagosome enriched proteins highlights a number of overrepresented biological processes including protein translation, protein localization and targeting, mRNA degradation and protein complex disassembly as well as ribosomal complexes. Our results show clear evidence for a causal relationship between NGLY1 disruption and induction of cellular autophagy, a result which may have implications for novel treatment modalities for NGLY1 disorder. Our findings furthermore make a strong case for the preferential use of Q-VD-OPh for cellular studies of caspase inhibition where an off-target mediated induction of autophagy is undesirable.

## Methods

### Methylthiazolyl diphenyl tetrazolium bromide (MTT) cell viability assay

Cells were plated in a 96 well plate (Greiner) at a density of 2000 cells per well. Cells were incubated with treatments as indicated per experiment. Following treatment, media was aspirated and replaced with fresh media (100 μl) containing MTT (final concentration 0.2 mg/ml). Plates were incubated at 37 °C for 2 h. Following incubation, media containing MTT was carefully removed and DMSO (100 μL) was added to each well. The plates were then shaken for 20 min at RT. The absorbance was read using a BMG Labtech FLUOstar OPTIMA plate reader at 570 nm. Treatment with menadione (20 μM, 24 h) was included as a positive control to confirm decreased absorbance in non-viable cells.

### Quantification of levels of GFP-LC3 positive puncta

GFP-LC3 HEK 293 cells were plated on uncoated 16 mm glass coverslips at (40 000 cells per well) and treated with Z-VAD-fmk or Q-VD-OPh (50 μM, 24-72 h) or a vehicle control. Cells were imaged on the oil immersion 63x objective. A minimum of three areas per coverslip were imaged and were selected under bright field. The number of GFP-LC3 positive puncta were counted manually and, the average number of puncta per cell was calculated per condition. To measure flux, GFP-LC3 HEK 293 cells were treated with Bafilomycin A1 (100 nM, 1 h) or 3-MA (5 mM, 1 h) at 37 °C at 5% CO2. Cells were re-imaged and the number of GFP-LC3 puncta was calculated.

### qPCR determination of ER stress markers

Assessment of ER stress markers was carried out by quantitative real-time PCR (qPCR). qPCR was used to determine changes in ER stress markers and *N*-glycanase downregulation for siRNA mediated knockdown. The sequences of all primers used are presented in Table 1. RT-qPCR was carried out using an MJ Opticom real time PCR machine. For analysis of ER stress markers, cells were treated with Z-VAD-fmk or Q-VD-OPh (50 μM, 24-72 h) or a vehicle control and MG132 (5 μM, 18 h). RNA was extracted using RNeasy Mini Kit. RNA (100 ng) was used with the One-Step Luna^®^ Universal qPCR Master Mix per manufacturer’s instructions. Samples were heated to 55 °C, 10 min to convert RNA to cDNA followed by an initial denaturation at 98°C, 5 min followed by denaturation at 95 °C, 30 s, annealing at 58 °C, 30s, extension at 72°C, 30 s for 42 cycles. The threshold was taken over the global minimum over 10 standard deviations and analysed using the double delta Ct method. GAPDH was employed as a housekeeping gene and an untreated control was used as the baseline.

### Propidium iodide exclusion and Annexin V staining

Cells were grown in DMEM media and plated into 12 well plate (Greiner). HEK 293 cells were treated with either Q-VD-OPh or Z-VAD-fmk (50 μM, 24-72 h) or a vehicle control (DMSO). Menadione (20 μM, 18 h) was used as a positive control for cell death and apoptosis induction. Samples were treated with trypsin to detach cells and washed three times in PBS. Cells were incubated with the Molecular Probes™ Dead Cell Apoptosis Kit with Annexin V FITC and PI, for flow cytometry. Cells were re-suspended in 1x annexin binding buffer (100 μl). FITC annexin V (5 μl) and Propidium iodide (PI) (1 μl, 100 μg/ml) were added for 15 min at RT. Samples were diluted with annexin binding buffer (400 μl) and red/green signal was analysed using a TALI image-based cytometer. The percentage of annexin V positive and PI negative cells were calculated, and cell number was recorded.

### Deglycosylation Dependent Venus (ddVENUS) assay

HEK 293 cells plated in 12 well plates (Greiner) at a density of 80 000 cells per well were transfected with ddVENUS plasmid DNA (2 μg per well) using JetPEI^®^ HTS DNA transfection reagent (Polyplus-transfection^®^) in accordance with the manufacturer protocol. After 24 h, Z-VAD-fmk (0-300 μM) was added and the cells incubated for a further 24 h. For genetic ablation of *N*-glycanase, HEK 293 cells were transfected with SMARTpool: ON-TARGETplus NGLY1 siRNA (25 nM) or an ON-TARGETplus non-targeting control (25 nM) using JetPRIME^®^ DNA and siRNA transfection reagent (Polyplus-transfection^®^) as per the manufacturer instructions. Cells were analysed 3d post-transfection.

The proteasome inhibitor, MG132 (8 μM) was added for 6 h before the cells were trypsinised, collected by centrifugation and re-suspended in HBSS (500 μl). Cells were kept on ice before measurement. Fluorescence intensity was measured by flow cytometry on a BD FACSCalibur (BD Biosciences) and analysed using BD CellQuest™. A minimum of 10 000 cells were analysed for each condition. A non-transfected well was used as a control to determine background fluorescence. Cells were analysed using the FL1 filter at 400 V. The non-transfected cells were gated out of analysis and the median fluorescence intensity of transfected cells was plotted.

### Intracellular calcium fluorescence imaging

GFP-LC3 HEK 293 cells were plated on uncoated 16 mm glass coverslips at 40 000 cells per well and treated with Z-VAD-fmk, Q-VD-OPh (50 μM, 24-72 h) or a vehicle control. For genetic ablation of *N*-glycanase, HEK 293 cells were transfected with SMARTpool: ON-TARGETplus NGLY1 siRNA (25 nM) or an ON-TARGETplus non-targeting control siRNA (25 nM) and analysed 3-5 d post-transfection. Cells were incubated with FURA-2 AM (1 μM) in imaging buffer (121 mM NaCl, 5.4 mM KCl, 0.8 mM MgCl_2_, 1.8 mM CaCl_2_, 6 mM NaHCO_3_, 5.5 mM D-glucose, 25 mM HEPES, pH 7.4 supplemented with Gibco MEM Amino Acids and MEM Non-Essential Amino Acids solution) for 20 min at 37°C. Cells were washed three times with imaging buffer and incubated for a further 20 min. Coverslips were imaged using a Leica DMI6000 fluorescence microscope on the 20x objective. Ca^2+^ was mobilized from intracellular stores using Thapsigargin (1 μM) after 180 s. Images were taken every 20 s for 12 m. Time lapse videos were analyzed using ImageJ. Cytosolic areas of cells were selected plus a background region for 340 and 380 nm. A minimum of 20 cellular regions were analyzed per coverslip. The 340/380 ratio was calculated. The area under the curve, baseline average of the first eight images and peak height were calculated using GraphPad Prism 7.

### Label-free Proteomics Analysis

GFP-LC3 HEK 293 cells were cultured as described above and treated with Z-VAD-fmk (50 μM) for 72 h or transfected with NGLY1 siRNA (25 nM) and used 5 days post-transfection. Cells (80 million per replicate) were treated with Bafilomycin A1 (100 nM) for 1 h to allow accumulation of autophagosomes. Following treatment with Bafilomycin, cells were washed twice with ice cold PBS and harvested by treatment with 0.25% Trypsin-EDTA (Thermofisher). Cells were pelleted and thrice washed in ice cold PBS. Cells were lysed in 1 % Triton in PBS by Dounce homogenization and incubated for 0.5 h at 4°C. The resultant lysate was centrifuged at 6000 g for 10 min at 4 °C. The supernatant was removed and centrifuged again at 20 000 g for 20 min at 4 °C. The supernatant was discarded, and the pellet re-suspended in PBS (500 μl) and added to filter spin columns (Corning) with GFPTrap^®^ agarose beads (50 μl) (Chromotek) or control agarose beads (50 μl) (Chromotek) and then washed three times in PBS. Samples were incubated with the agarose beads and tumbled for 1 h at 4°C. The spin filter columns were centrifuged at 2 500 g for 2 min and, the flow-through discarded. Agarose beads were washed 7x in ice-cold PBS-T by centrifugation. Samples were eluted with 0.2 M glycine, pH 2.5 (50 μl) for 1 min with constant vortexing. Beads were centrifuged into a fresh, protein Lobind tubes (Eppendorf). Elution was repeated and the eluates were pooled. Samples were neutralized by the addition of 1 M phosphate buffer (pH 7.4). Eluted material was then processed for proteomics analysis by filter-aided sample preparation (FASP) (Wiśniewski et al., 2009). Tryptic digests were desalted and then analyzed by liquid chromatography tandem mass spectrometry (LC-MS/MS). Raw data were processed in the MaxQuant software utilizing the MaxLFQ label-free quantification algorithm. Processed data were further analyzed using the Perseus computational platform. After removal of known contaminant proteins and requiring quantification in at least two out of three biological replicates in at least one group, a total of 915 protein hits were obtained. Missing values were imputed (replacement of missing values from normal distribution; width 0.5, down shift 1.4) as described previously (Keilhauer et al., 2015). Protein enrichment and depletion were visualized using volcano plots and heatmap with hierarchical clustering analysis (HCA). Principal component analysis (PCA) was used to visualize protein abundance relationships across all samples. A schematic of the workflow is presented in (Fig S5a).

## Accession Numbers

The mass spectrometry proteomics data have been deposited to the ProteomeXchange Consortium via the PRIDE (Perez-Riverol et al., 2019) partner repository with the dataset identifier PXD020367.

## Acknowledgements

SHN was supported through funding provided by The Open University; HBK was supported by the Wellcome Trust OXION Ion Channels Initiative. Atg13^-/-^ MEFs and corresponding WT MEFs were a kind gift from Nick Ktistakis, Babraham Institute.

## Author Contributions

SHN: methodology, formal analysis, investigation, data curation, writing – original draft. MDB: supervision, writing – review and editing. HBK: supervision, writing – review and editing, mass spectrometry, proteomics data analysis. JEG: resources, writing – review and editing. SAA: conceptualization, supervision, writing – original draft.

---

**Table 1.**
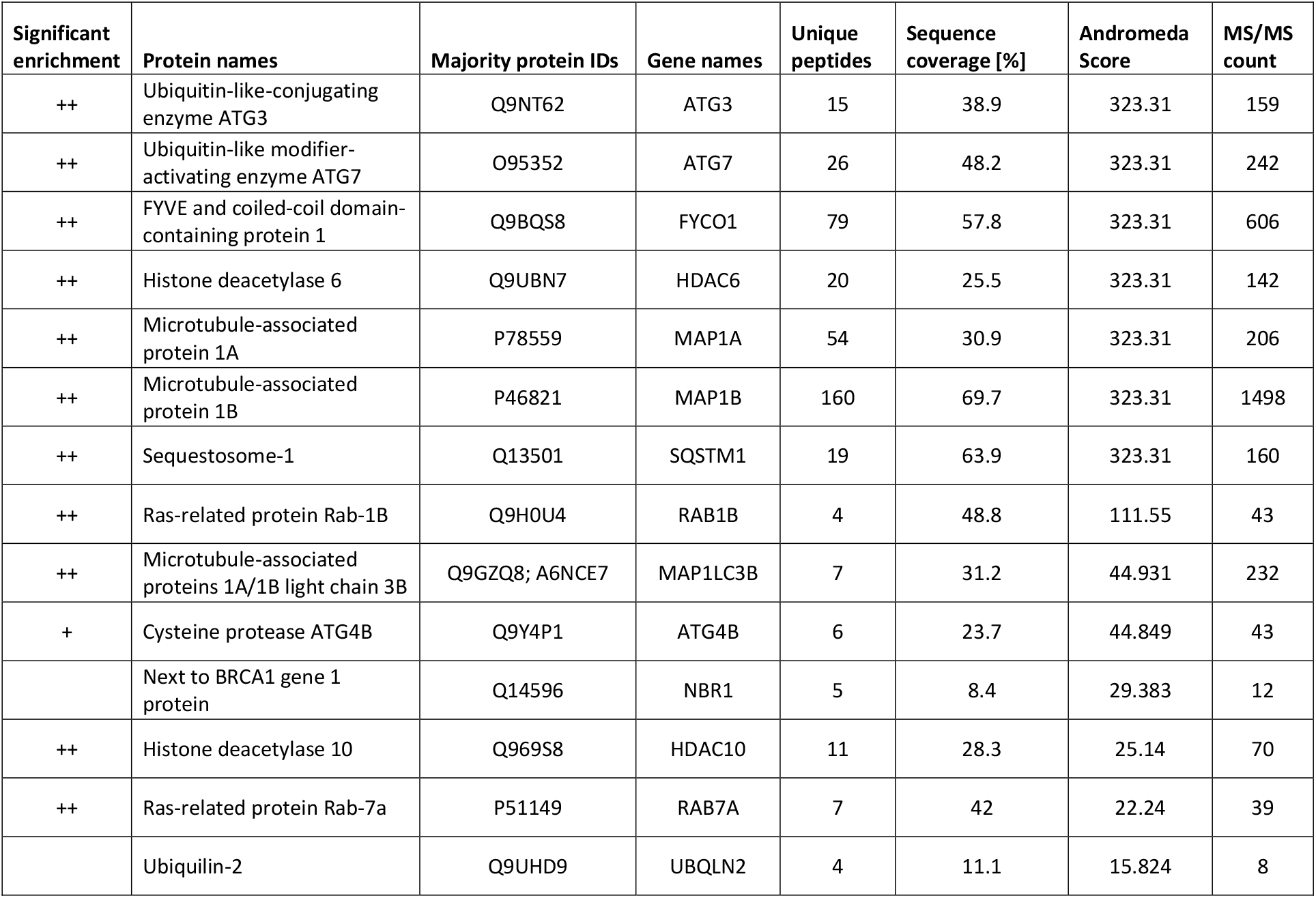
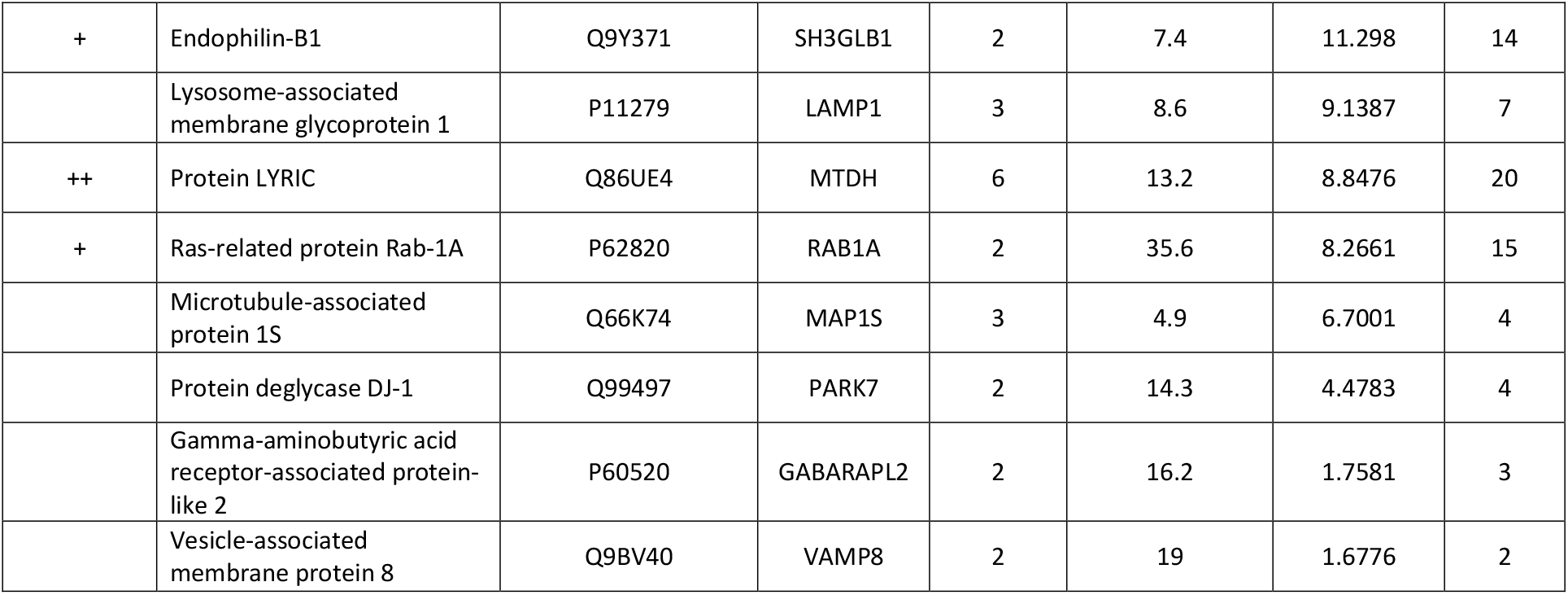
Enrichment of autophagy-related proteins in autophagosome IPs following Z-VAD-fmk or NGLY1 KD. Tabulated autophagy-related proteins shown with number of unique peptides identified, percentage sequence coverage, Andromeda score and MS/MS count. Only protein hits with ≥ 2 tryptic peptides/protein are displayed. Significant enrichment: ‘++’ significantly enriched in both autophagosome IPs; ‘+’ significantly enriched in one autophagosome IP (either following Z-VAD-fmk treatment or NGLY1 KD); ‘ ‘ no significant enrichment in either autophagosome IP.

| Significant enrichment | Protein names | Majority protein IDs | Gene names | Unique peptides | Sequence coverage [%] | Andromeda Score | MS/MS count |
| --- | --- | --- | --- | --- | --- | --- | --- |
| ++ | Ubiquitin-like-conjugating enzyme ATG3 | Q9NT62 | ATG3 | 15 | 38.9 | 323.31 | 159 |
| ++ | Ubiquitin-like modifier-activating enzyme ATG7 | O95352 | ATG7 | 26 | 48.2 | 323.31 | 242 |
| ++ | FYVE and coiled-coil domain-containing protein 1 | Q9BQS8 | FYCO1 | 79 | 57.8 | 323.31 | 606 |
| ++ | Histone deacetylase 6 | Q9UBN7 | HDAC6 | 20 | 25.5 | 323.31 | 142 |
| ++ | Microtubule-associated protein 1A | P78559 | MAP1A | 54 | 30.9 | 323.31 | 206 |
| ++ | Microtubule-associated protein 1B | P46821 | MAP1B | 160 | 69.7 | 323.31 | 1498 |
| ++ | Sequestosome-1 | Q13501 | SQSTM1 | 19 | 63.9 | 323.31 | 160 |
| ++ | Ras-related protein Rab-1B | Q9H0U4 | RAB1B | 4 | 48.8 | 111.55 | 43 |
| ++ | Microtubule-associated proteins 1A/1B light chain 3B | Q9GZQ8; A6NCE7 | MAP1LC3B | 7 | 31.2 | 44.931 | 232 |
| + | Cysteine protease ATG4B | Q9Y4P1 | ATG4B | 6 | 23.7 | 44.849 | 43 |
|  | Next to BRCA1 gene 1 protein | Q14596 | NBR1 | 5 | 8.4 | 29.383 | 12 |
| ++ | Histone deacetylase 10 | Q969S8 | HDAC10 | 11 | 28.3 | 25.14 | 70 |
| ++ | Ras-related protein Rab-7a | P51149 | RAB7A | 7 | 42 | 22.24 | 39 |
|  | Ubiquilin-2 | Q9UHD9 | UBQLN2 | 4 | 11.1 | 15.824 | 8 |
| + | Endophilin-B1 | Q9Y371 | SH3GLB1 | 2 | 7.4 | 11.298 | 14 |
|  | Lysosome-associated<br>membrane glycoprotein 1 | P11279 | LAMP1 | 3 | 8.6 | 9.1387 | 7 |
| ++ | Protein LYRIC | Q86UE4 | MTDH | 6 | 13.2 | 8.8476 | 20 |
| + | Ras-related protein Rab-1A | P62820 | RAB1A | 2 | 35.6 | 8.2661 | 15 |
|  | Microtubule-associated<br>protein 1S | Q66K74 | MAP1S | 3 | 4.9 | 6.7001 | 4 |
|  | Protein deglycase DJ-1 | Q99497 | PARK7 | 2 | 14.3 | 4.4783 | 4 |
|  | Gamma-aminobutyric acid<br>receptor-associated protein-<br>like 2 | P60520 | GABARAPL2 | 2 | 16.2 | 1.7581 | 3 |
|  | Vesicle-associated<br>membrane protein 8 | Q9BV40 | VAMP8 | 2 | 19 | 1.6776 | 2 |

**Supplementary Figure S1.**
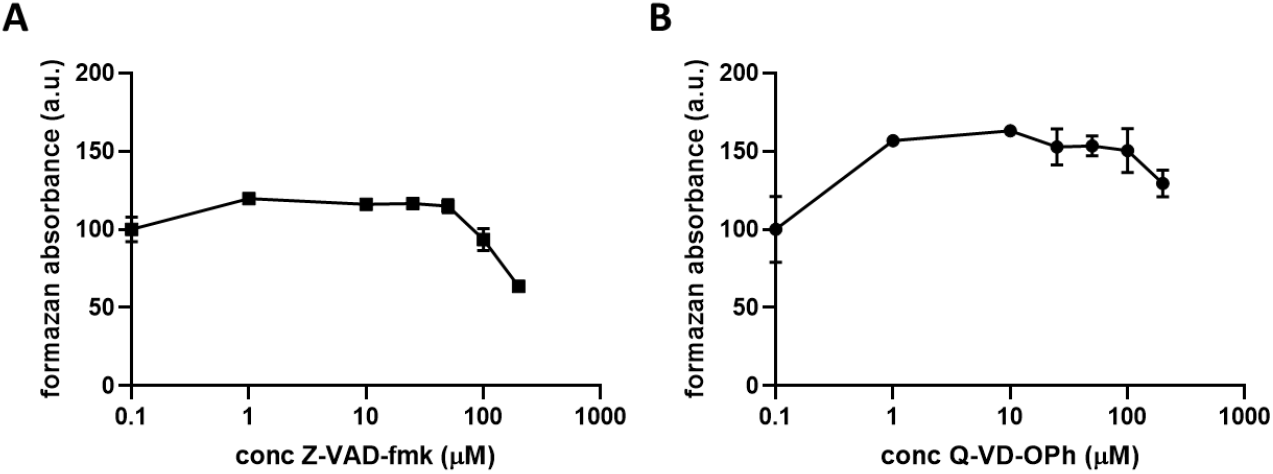
Z-VAD-fmk is not toxic under 200 μM. MTT assay of HEK 293 cells were treated with Z-VAD-fmk or Q-VD-OPh for 24 h. Data was normalised to the vehicle. One-way ANOVA, followed by a Dunnett’s post hoc test against the vehicle. n=3, * indicates P < 0.05, error bars ± SEM.

**Supplementary Figure S2.**
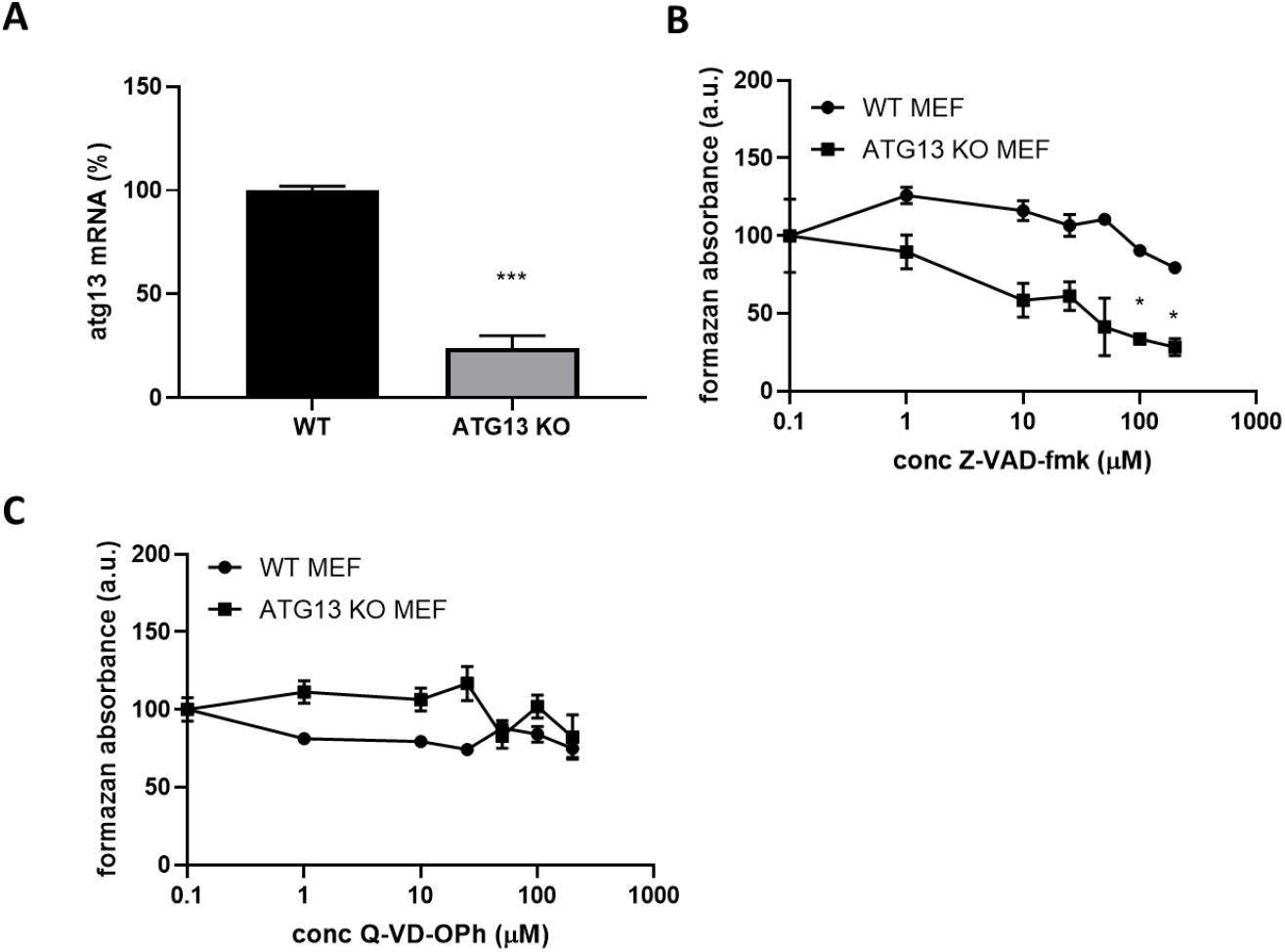
Z-VAD-fmk reduced the ability of autophagy deficient cells to reduce MTT. **(A)** MTT of ATG13 KO MEF and WT MEF treated with (**B**) Z-VAD-fmk (1-200 μM) or (**C**) Q-VD-OPh (1-200 μM) for 24 h. Multiple t-tests, Holm-Sidak. Error bars ± SEM, n=3, p < 0.05.

**Supplementary Figure S3.**
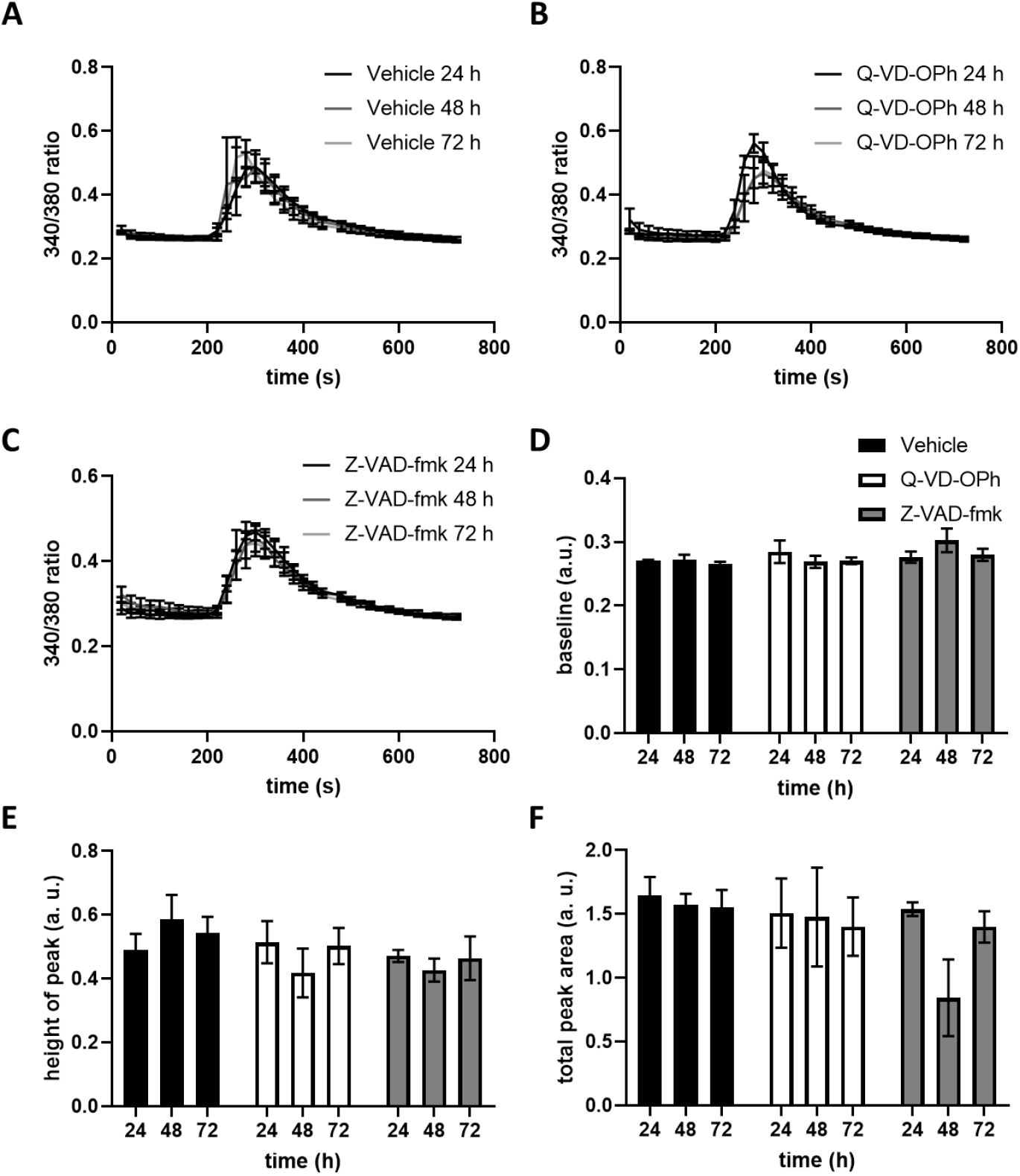
Z-VAD-fmk or Q-VD-OPh treatment does not affect Ca^2+^ handling. (**A-C**) 340/380 ratio traces of treated (vehicle, Q-VD-OPh or Z-VAD-fmk) HEK 293 cells loaded with Fura-2 AM and treated with thapsigargin (1 μM) at 180 s. (**D**) The baseline was averaged for the first 100 s (**E**) the maximum peak height was calculated and (**F**) area under the curve was determined. n=3, error bars indicate ± SEM.

**Supplementary Figure S4.**
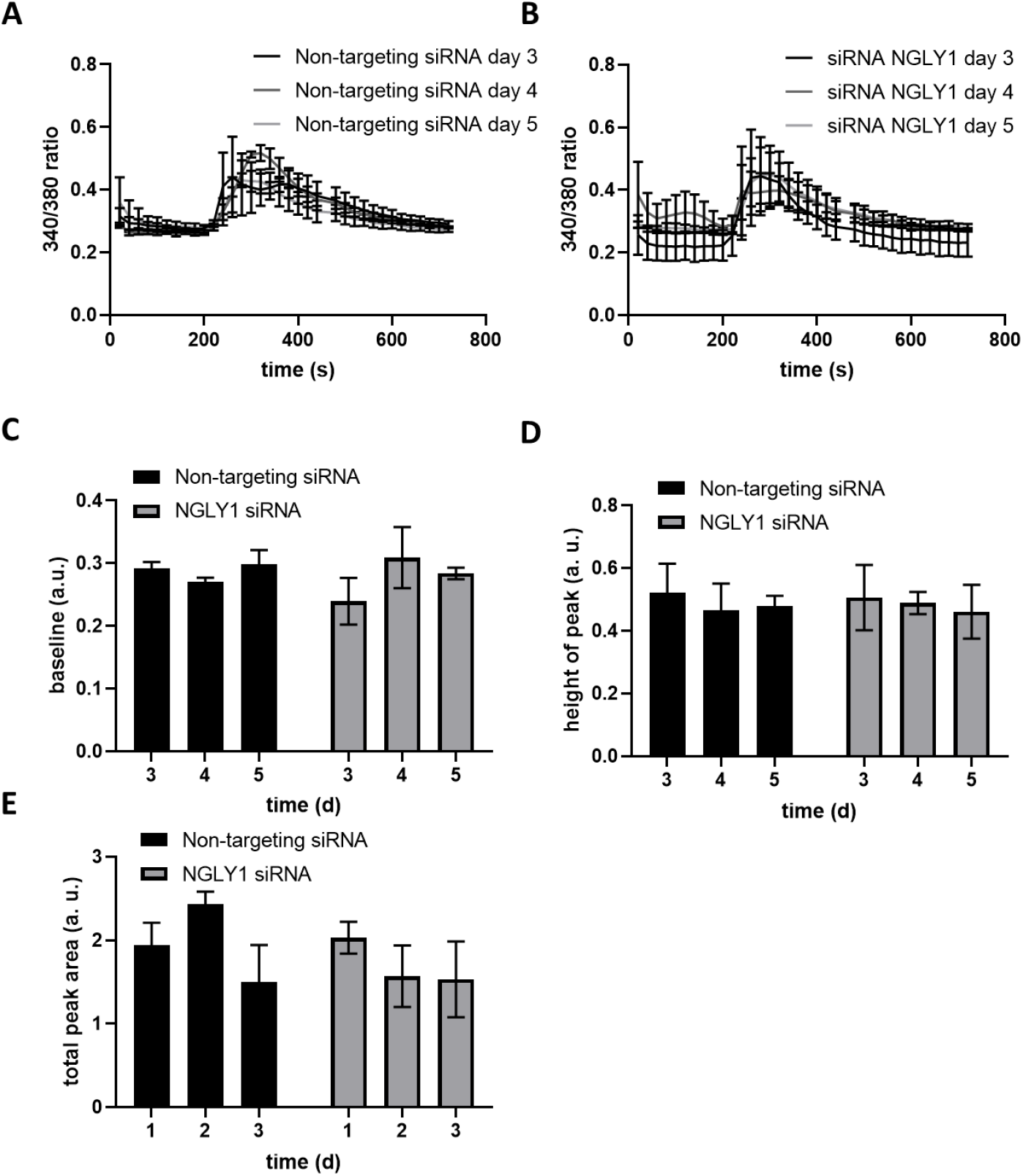
NGLY1 siRNA knockdown does not affect Ca^2+^ handling. (**A-B**) 340/380 ratio traces of HEK 293 cells (transfected with NGLY1 siRNA or non-targeting siRNA) loaded with Fura-2 AM and treated with thapsigargin (1 μM) at 180 s. (**C**) The baseline was averaged for the first 100 s, (D) the maximum peak height was calculated and (**E**) the area under the curve was taken. n=3, error bars indicate ± SEM.

**Supplementary Figure S5.**
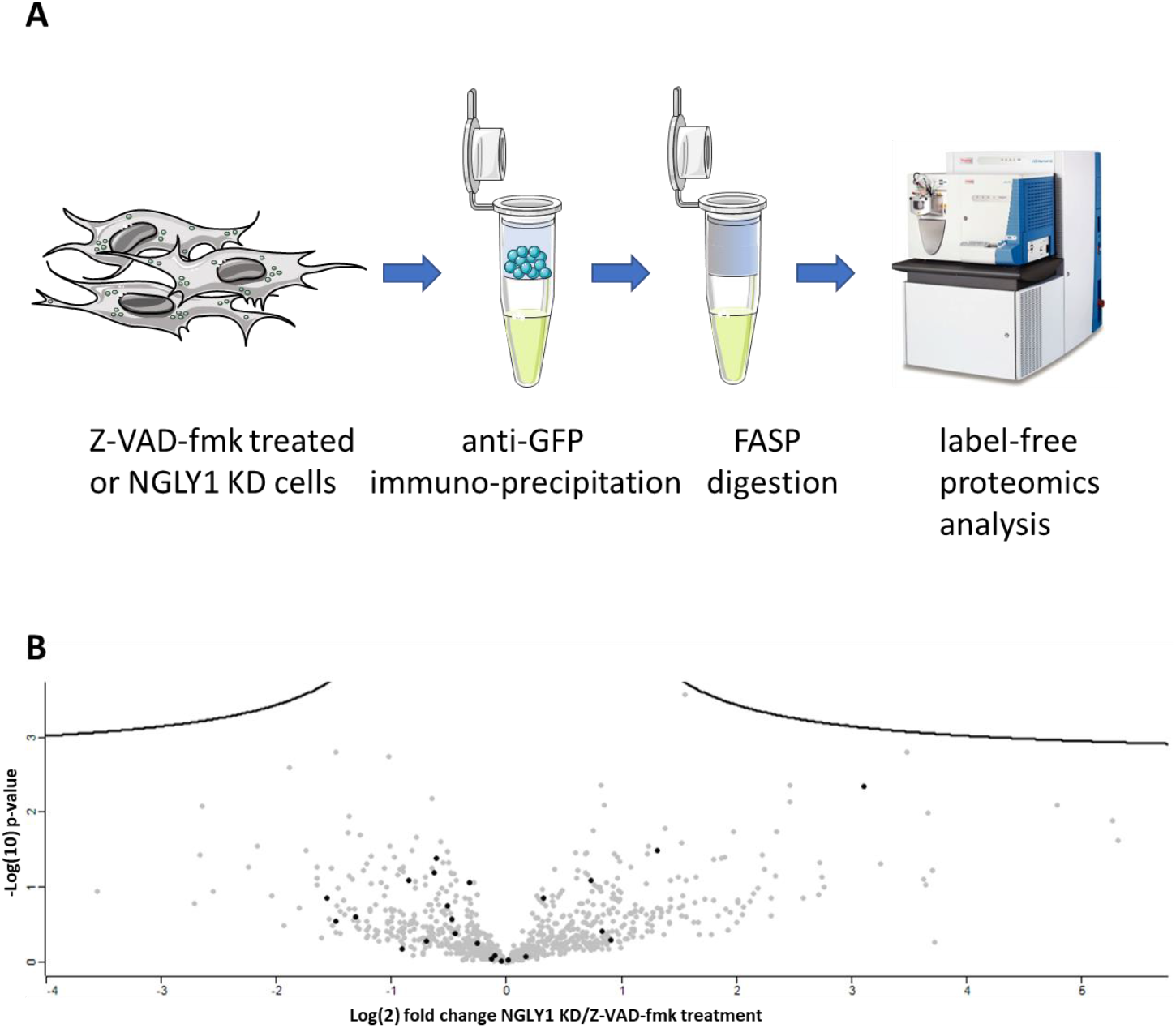
Schematic representation of proteomics analysis workflow. **(A)** Schematic representation of workflow for label-free proteomics experiments (vector graphics adapted from Servier SMART Medical Art (https://smart.servier.com/) and used here under the terms of Creative Commons Attribution 3.0 Unported License **(B)** Label-free proteomics analysis of autophagosomes shows no significantly changing proteins between NGLY1 KD and Z-VAD-fmk treatment; Volcano plot (FDR 0.05, s0 0.1) showing autophagy-related protein hits not significantly altered (black).

**Supplementary Figure S6.**
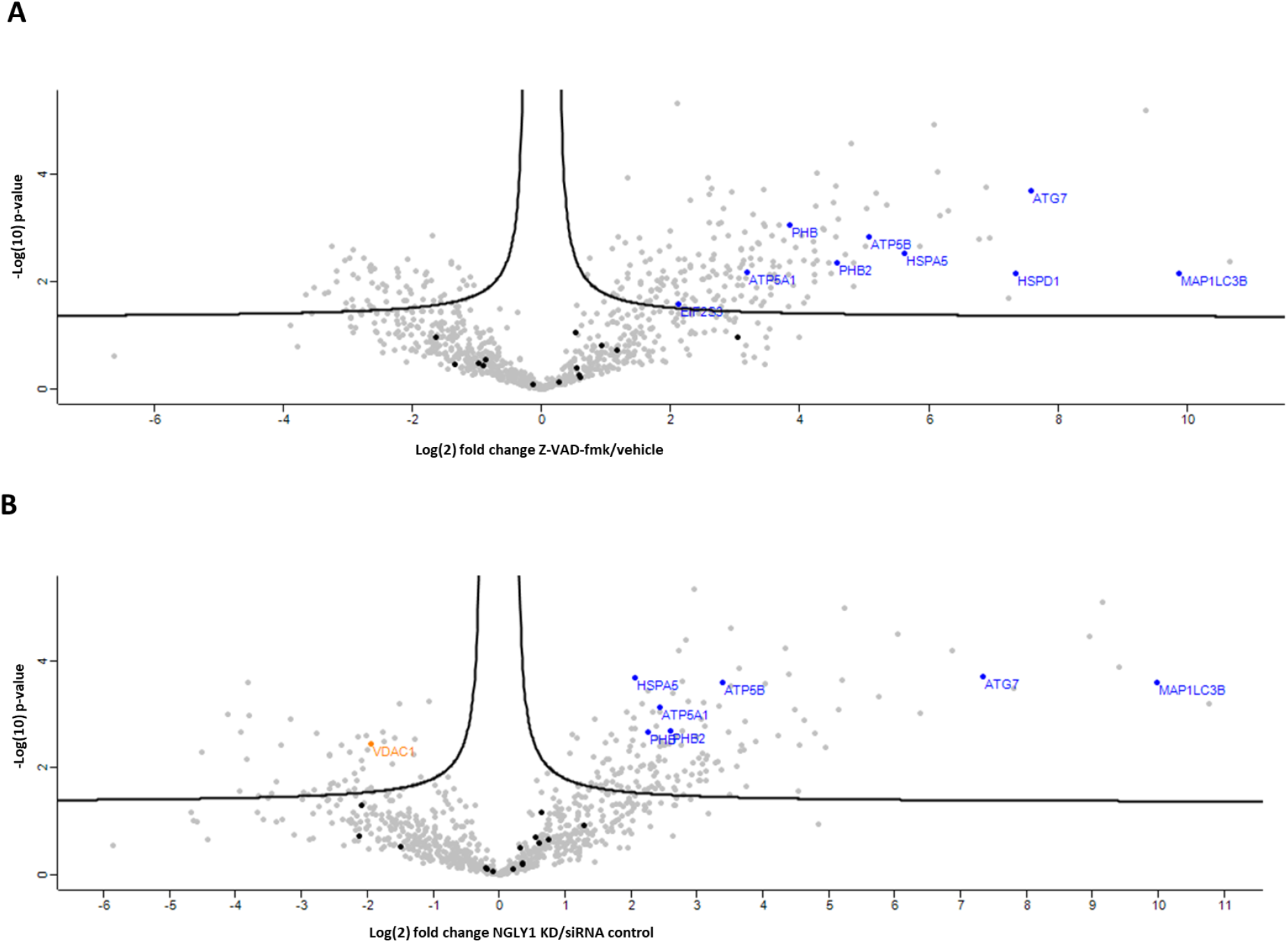
Comparison to: Gao, W., *et al., Biochemical isolation and characterization of the tubulovesicular LC3-positive autophagosomal compartment*. Journal of Biological Chemistry, 2010. 285(2): p. 1371-1383. **(A)** Z-VAD-fmk treatment: 23 out of 101 proteins identified; of these 9 significantly enriched (blue) and 14 not significantly altered (black) **(B)** NGLY1 KD 23 out of 101 proteins identified; of these 7 significantly enriched (blue), 1 significantly depleted (orange) and 15 not significantly altered (black); Volcano plot (FDR 0.05, s0 0.1).

**Supplementary Figure S7.**
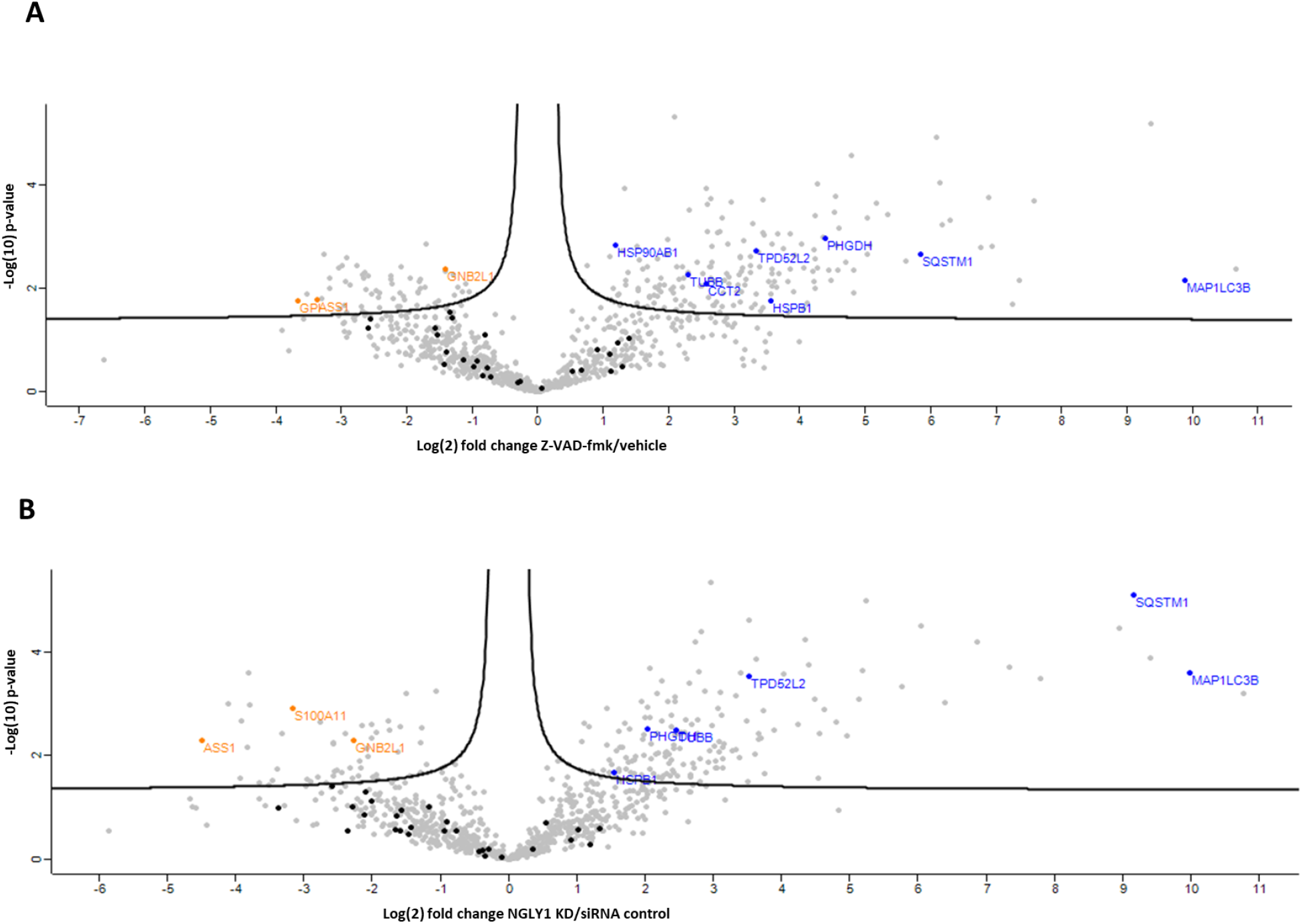
Comparison to: Dengjel, J., *et al., Identification of autophagosome-associated proteins and regulators by quantitative proteomic analysis and genetic screens*. Molecular and Cellular Proteomics, 2012. 11(3):M111.014035. **(A)** Z-VAD-fmk treatment: 37 out of 94 proteins were identified in our analysis; of these 8 significantly enriched (blue), 3 significantly depleted (orange) and 26 not significantly altered (black) **(B)** NGLY1 KD 37 out of 94 proteins identified; of these 6 significantly enriched (blue), 3 significantly depleted (orange) and 28 not significantly altered (black); Volcano plot (FDR 0.05, s0 0.1).

**Supplementary Table ST1. Gene ontology enrichment analysis - hierarchical clustering analysis (HCA) cluster I – protein hits enriched in autophagosome IPs** Significantly overrepresented (Benjamini-Hochberg FDR <0.05) Gene Ontology (GO) terms for biological processes (GOBP), cellular compartment (GOCC) and molecular function (GOMF); total number of protein hits in cluster II: 228

**Supplementary Table ST2. Gene ontology enrichment analysis - hierarchical clustering analysis (HCA) cluster II – protein hits enriched in negative control IPs** Significantly overrepresented (Benjamini-Hochberg FDR <0.05) Gene Ontology (GO) terms for biological processes (GOBP), cellular compartment (GOCC) and molecular function (GOMF); total number of protein hits in cluster II: 141

## Highlights

- Autophagy is induced both on treatment of cells with Z-VAD-fmk and in NGLY1 KD
- Autophagic flux is not impaired by Z-VAD-fmk treatment or NGLY1 KD in HEK293
- Autophagosomal protein content is comparable in Z-VAD-fmk treatment and NGLY1 KD
- Caspase inhibitor Q-VD-OPh does not exhibit NGLY1-mediated off-target effects

## References

Asahina, M., Fujinawa, R., Nakamura, S., Yokoyama, K., Tozawa, R., and Suzuki, T. (2020). Ngly1 -/- rats develop neurodegenerative phenotypes and pathological abnormalities in their peripheral and central nervous systems. Human Molecular Genetics 29, 1635–1647.

Bhutia, S.K., Kegelman, T.P., Das, S.K., Azab, B., Su, Z.Z., Lee, S.G., Sarkar, D., and Fisher, P.B. (2010). Astrocyte elevated gene-1 induces protective autophagy. Proceedings of the National Academy of Sciences of the United States of America 107, 22243–22248.

Caserta, T.M., Smith, A.N., Gultice, A.D., Reedy, M.A., and Brown, T.L. (2003). Q-VD-OPh, a broad spectrum caspase inhibitor with potent antiapoptotic properties. Apoptosis 8, 345–352.

Chauvier, D., Ankri, S., Charriaut-Marlangue, C., Casimir, R., and Jacotot, E. (2007). Broad-spectrum caspase inhibitors: From myth to reality? [5]. Cell Death and Differentiation 14, 387–391.

Chen, S.Y., Chiu, L.Y., Maa, M.C., Wang, J.S., Chien, C.L., and Lin, W.W. (2011). zVAD-induced autophagic cell death requires c-Src-dependent ERK and JNK activation and reactive oxygen species generation. Autophagy 7, 217–228.

Cox, J., Hein, M.Y., Luber, C.A., Paron, I., Nagaraj, N., and Mann, M. (2014). Accurate proteome-wide label-free quantification by delayed normalization and maximal peptide ratio extraction, termed MaxLFQ. Molecular and Cellular Proteomics 13, 2513–2526.

Cox, J., and Mann, M. (2008). MaxQuant enables high peptide identification rates, individualized p.p.b.-range mass accuracies and proteome-wide protein quantification. Nature Biotechnology 26, 1367–1372.

Dengjel, J., Høyer-Hansen, M., Nielsen, M.O., Eisenberg, T., Harder, L.M., Schandorff, S., Farkas, T., Kirkegaard, T., Becker, A.C., Schroeder, S., et al. (2012). Identification of autophagosome-associated proteins and regulators by quantitative proteomic analysis and genetic screens. Molecular and Cellular Proteomics 11.

Deszcz, L., Seipelt, J., Vassilieva, E., Roetzer, A., and Kuechler, E. (2004). Antiviral activity of caspase inhibitors: Effect on picornaviral 2A proteinase. FEBS Letters 560, 51–55.

Eichhold, T.H., Hookfin, E.B., Taiwo, Y.O., De, B., and Wehmeyer, K.R. (1997). Isolation and quantification of fluoroacetate in rat tissues, following dosing of 2-Phe-Ala-CH2-F, a peptidyl fluoromethyl ketone protease inhibitor. Journal of Pharmaceutical and Biomedical Analysis 16, 459–467.

Enns, G.M., Shashi, V., Bainbridge, M., Gambello, M.J., Zahir, F.R., Bast, T., Crimian, R., Schoch, K., Platt, J., Cox, R., et al. (2014). Mutations in NGLY1 cause an inherited disorder of the endoplasmic reticulum-associated degradation pathway. Genetics in medicine : official journal of the American College of Medical Genetics 16, 751–758.

Fujihira, H., Masahara-Negishi, Y., Tamura, M., Huang, C., Harada, Y., Wakana, S., Takakura, D., Kawasaki, N., Taniguchi, N., Kondoh, G., et al. (2017). Lethality of mice bearing a knockout of the Ngly1-gene is partially rescued by the additional deletion of the Engase gene. PLoS Genetics 13.

Gao, W., Kang, J.H., Liao, Y., Ding, W.X., Gambotto, A.A., Watkins, S.C., Liu, Y.L., Stolz, D.B., and Yin, X.M. (2010). Biochemical isolation and characterization of the tubulovesicular LC3-positive autophagosomal compartment. Journal of Biological Chemistry 285, 1371–1383.

Garcia-Calvo, M., Peterson, E.P., Leiting, B., Ruel, R., Nicholson, D.W., and Thornberry, N.A. (1998). Inhibition of human caspases by peptide-based and macromolecular inhibitors. Journal of Biological Chemistry 273, 32608–32613.

Grotzke, J.E., Lu, Q., and Cresswell, P. (2013). Deglycosylation-dependent fluorescent proteins provide unique tools for the study of ER-associated degradation. Proceedings of the National Academy of Sciences of the United States of America 110, 3393–3398.

Harrison, B., Kraus, M., Burch, L., Stevens, C., Craig, A., Gordon-Weeks, P., and Hupp, T.R. (2008). DAPK-1 binding to a linear peptide motif in MAP1B stimulates autophagy and membrane blebbing. Journal of Biological Chemistry 283, 9999–10014.

He, P., Grotzke, J.E., Ng, B.G., Gunel, M., Jafar-Nejad, H., Cresswell, P., Enns, G.M., and Freeze, H.H. (2015). A congenital disorder of deglycosylation: Biochemical characterization of N-glycanase 1 deficiency in patient fibroblasts. Glycobiology 25, 836–844.

Kabeya, Y., Mizushima, N., Ueno, T., Yamamoto, A., Kirisako, T., Noda, T., Kominami, E., Ohsumi, Y., and Yoshimori, T. (2000). LC3, a mammalian homologue of yeast Apg8p, is localized in autophagosome membranes after processing. EMBO Journal 19, 5720–5728.

Kaizuka, T., and Mizushima, N. (2016). Atg13 is essential for autophagy and cardiac development in mice. Molecular and Cellular Biology 36, 585–595.

Keilhauer, E.C., Hein, M.Y., and Mann, M. (2015). Accurate protein complex retrieval by affinity enrichment mass spectrometry (AE-MS) rather than affinity purification mass spectrometry (AP-MS). Molecular and Cellular Proteomics 14, 120–135.

Kim, R.J., Hah, Y.S., Sung, C.M., Kang, J.R., and Park, H.B. (2014). Do antioxidants inhibit oxidative-stress-induced autophagy of tenofibroblasts? Journal of Orthopaedic Research 32, 937–943.

Kong, J., Peng, M., Ostrovsky, J., Kwon, Y.J., Oretsky, O., McCormick, E.M., He, M., Argon, Y., and Falk, M.J. (2018). Mitochondrial function requires NGLY1. Mitochondrion 38, 6–16.

Lee, J.Y., Koga, H., Kawaguchi, Y., Tang, W., Wong, E., Gao, Y.S., Pandey, U.B., Kaushik, S., Tresse, E., Lu, J., et al. (2010). HDAC6 controls autophagosome maturation essential for ubiquitin-selective quality-control autophagy. EMBO Journal 29, 969–980.

Lehrbach, N.J., Breen, P.C., and Ruvkun, G. (2019). Protein Sequence Editing of SKN-1A/Nrf1 by Peptide:N-Glycanase Controls Proteasome Gene Expression. Cell 177, 737–750.e715.

Luyten, T., Welkenhuyzen, K., Roest, G., Kania, E., Wang, L., Bittremieux, M., Yule, D.I., Parys, J.B., and Bultynck, G. (2017). Resveratrol-induced autophagy is dependent on IP3Rs and on cytosolic Ca2 +. Biochimica et Biophysica Acta - Molecular Cell Research 1864, 947–956.

Mann, S.S., and Hammarback, J.A. (1994). Molecular characterization of light chain 3. A microtubule binding subunit of MAP1A and MAP1B. Journal of Biological Chemistry 269, 11492–11497.

Maynard, J.C., Fujihira, H., Dolgonos, G.E., Suzuki, T., and Burlingame, A.L. (2020). Cytosolic N-GlcNAc proteins are formed by the action of endo-β-N-acetylglucosaminidase. Biochemical and Biophysical Research Communications 530, 719–724.

Misaghi, S., Pacold, M.E., Blom, D., Ploegh, H.L., and Korbel, G.A. (2004). Using a small molecule inhibitor of peptide: N-glycanaseto probe its role in glycoprotein turnover. Chemistry and Biology 11, 1677–1687.

Morrison, J.F., and Peters, R.A. (1954). Biochemistry of fluoroacetate poisoning: the effect of fluorocitrate on purified aconitase. The Biochemical journal 58, 473–479.

Mueller, W.F., Jakob, P., Sun, H., Clauder-Münster, S., Ghidelli-Disse, S., Ordonez, D., Boesche, M., Bantscheff, M., Collier, P., Haase, B., et al. (2020). Loss of N-glycanase 1 alters transcriptional and translational regulation in K562 cell lines. G3: Genes, Genomes, Genetics 10, 1585–1597.

Oehme, I., Linke, J.P., Böck, B.C., Milde, T., Lodrini, M., Hartenstein, B., Wiegand, I., Eckert, C., Roth, W., Kool, M., et al. (2013). Histone deacetylase 10 promotes autophagy-mediated cell survival. Proceedings of the National Academy of Sciences of the United States of America 110, E2592–E2601.

Owings, K.G., Lowry, J.B., Bi, Y., Might, M., and Chow, C.Y. (2018). Transcriptome and functional analysis in a Drosophila model of NGLY1 deficiency provides insight into therapeutic approaches. Human Molecular Genetics 27, 1055–1066.

Pankiv, S., Alemu, E.A., Brech, A., Bruun, J.A., Lamark, T., Øvervatn, A., Bjørkøy, G., and Johansen, T. (2010). FYCO1 is a Rab7 effector that binds to LC3 and PI3P to mediate microtubule plus end - Directed vesicle transport. Journal of Cell Biology 188, 253–269.

Perez-Riverol, Y., Csordas, A., Bai, J., Bernal-Llinares, M., Hewapathirana, S., Kundu, D.J., Inuganti, A., Griss, J., Mayer, G., Eisenacher, M., et al. (2019). The PRIDE database and related tools and resources in 2019: Improving support for quantification data. Nucleic Acids Research 47, D442–D450.

Rodriguez, T.P., Mast, J.D., Hartl, T., Lee, T., Sand, P., and Perlstein, E.O. (2018). Defects in the neuroendocrine axis contribute to global development delay in a Drosophila model of NGLY1 deficiency. G3: Genes, Genomes, Genetics 8, 2193–2204.

Schotte, P., Declercq, W., Van Huffel, S., Vandenabeele, P., and Beyaert, R. (1999). Non-specific effects of methyl ketone peptide inhibitors of caspases. FEBS Letters 442, 117–121.

Suzuki, T., Huang, C., and Fujihira, H. (2016). The cytoplasmic peptide: N-glycanase (NGLY1) - Structure, expression and cellular functions. Gene 577, 1–7.

Suzuki, T., Huang, C., Harada, Y., Hosomi, A., Masahara-Negishi, Y., Seino, J., Fujihira, H., Funakoshi, Y., Suzuki, T., and Dohmae, N. (2015). Endo-β-n-acetylglucosaminidase forms N-GlcNAc protein aggregates during ER-associated degradation in NGLY1-defective cells. Proceedings of the National Academy of Sciences of the United States of America 112, 1398–1403.

Suzuki, T., Seko, A., Kitajima, K., Inoue, Y., and Inoue, S. (1993). Identification of Peptide:N-Glycanase Activity in Mammalian-Derived Cultured Cells. Biochemical and Biophysical Research Communications 194, 1124–1130.

Tambe, M.A., Ng, B.G., and Freeze, H.H. (2019). N-Glycanase 1 Transcriptionally Regulates Aquaporins Independent of Its Enzymatic Activity. Cell Reports 29, 4620–4631.e4624.

Tang, Z., Lin, M.G., Stowe, T.R., Chen, S., Zhu, M., Stearns, T., Franco, B., and Zhong, Q. (2013). Autophagy promotes primary ciliogenesis by removing OFD1 from centriolar satellites. Nature 502, 254–257.

Tomlin, F.M., Gerling-Driessen, U.I.M., Liu, Y.C., Flynn, R.A., Vangala, J.R., Lentz, C.S., Clauder-Muenster, S., Jakob, P., Mueller, W.F., Ordoñez-Rueda, D., et al. (2017). Inhibition of NGLY1 Inactivates the Transcription Factor Nrf1 and Potentiates Proteasome Inhibitor Cytotoxicity. ACS Central Science 3, 1143–1155.

Tyanova, S., Temu, T., Sinitcyn, P., Carlson, A., Hein, M.Y., Geiger, T., Mann, M., and Cox, J. (2016). The Perseus computational platform for comprehensive analysis of (prote)omics data. Nature Methods 13, 731–740.

Waterhouse, N.J., Finucane, D.M., Green, D.R., Elce, J.S., Kumar, S., Alnemri, E.S., Litwack, G., Khanna, K., Lavin, M.F., and Watters, D.J. (1998). Calpain activation is upstream of caspases in radiation-induced apoptosis. Cell Death and Differentiation 5, 1051–1061.

Wiśniewski, J.R., Zougman, A., Nagaraj, N., and Mann, M. (2009). Universal sample preparation method for proteome analysis. Nature Methods 6, 359–362.

Wu, T., Li, Y., Huang, D., Han, F., Zhang, Y.Y., Zhang, D.W., and Han, J. (2014). Regulator of G-protein signaling 19 (RGS19) and its partner Gα-inhibiting activity polypeptide 3 (GNAI3) are required for zVAD-induced autophagy and cell death in L929 cells. PLoS ONE 9.

Wu, Y.T., Tan, H.L., Huang, Q., Sun, X.J., Zhu, X., and Shen, H.M. (2011). ZVAD-induced necroptosis in L929 cells depends on autocrine production of TNFα mediated by the PKC-MAPKs-AP-1 pathway. Cell Death and Differentiation 18, 26–37.

Yang, K., Huang, R., Fujihira, H., Suzuki, T., and Yan, N. (2018). N-glycanase NGLY1 regulates mitochondrial homeostasis and inflammation through NRF1. The Journal of experimental medicine 215, 2600–2616.

Zhang, J., Chiu, J., Zhang, H., Qi, T., Tang, Q., Ma, K., Lu, H., and Li, G. (2013). Autophagic cell death induced by resveratrol depends on the Ca2+/AMPK/mTOR pathway in A549 cells. Biochemical Pharmacology 86, 317–328.

